# Infection risk and extensive parental care govern the molecular evolution of *Toll-like* receptors in birds

**DOI:** 10.1101/860114

**Authors:** Zhechun Zhang, Dan Liang, Guoling Chen, Fasheng Zou, Fumin Lei, Lu Dong, Michael Griesser, Yang Liu

**Author notes:** These authors contributed equally (Zhechun Zhang, Dan Liang). Senior authors (Michael Griesser, Yang Liu).

## Abstract

The arms race between pathogens and the immune system of their hosts is a critical evolutionary force that affects the ecology and life history of organisms. An increased infection risk selects for adaptations in immune genes that encode receptors involved in pathogen recognition and the initiation of innate immune responses, including Toll-like receptor (*TLR*) genes. Although recent studies assessed the evolution of major histocompatibility complex (MHC) genes, the ecological and evolutionary processes that drive the evolution of immune genes across major phylogenetic lineages remain unstudied. Moreover, trade-offs between immune responses and other energy-demanding vital functions may limit the resource allocation into costly immune functions, and therefore affect the evolution of immune genes. Here we assess the evolutionary patterns of six *TLR* genes across 121 bird species, covering 95% of extant orders that represent diverse ecologies and life histories. Selection analyses revealed that all six *TLR* genes show strong signs of purifying selection, while few sites underwent episodic positive selection. Comparative phylogenetic analyses showed that the intensity of positive selection of *TLR* genes is associated with long-distance migration, extensive parental care (i.e., altricality and prolonged parent-offspring association), and a large body size (a proxy of increased longevity). Together, these results suggest that the evolution of immune genes is characterized by episodic positive selection, and is shaped by an increased inflection risk and extensive parental care that buffers the costs of immune functions.

## Introduction

The immune system is critical for the survival of organisms as it mitigates the harmful effects of parasites and pathogens (bacteria, protozoa, and viruses). In response to the immune response of their hosts, pathogens continuously evolve, and understanding this co-evolutionary arms race is central for evolutionary biology (Ozinsky, et al. 2000; Hedrick 2002; Kumar, et al. 2011). Although quantifying selective pressures on host species is challenging in natural populations, it is possible to investigate pathogen-mediated selection on immune genes involved in the interaction with pathogen antigens (Hedrick 2002; Woolhouse, et al. 2002). Studies on of immune genes focus particularly on major histocompatibility complex (MHC) genes that govern acquired immune responses (Hedrick 2002; Bernatchez and Landry 2003; Acevedo-Whitehouse and Cunningham 2006), and Toll-like receptors (*TLR*) genes that activate innate immune responses (Sheldon and Verhulst 1996; Akira and Takeda 2004; Acevedo-Whitehouse and Cunningham 2006). Studies showed that functional constraints favor purifying selection of immune genes, but some studies did also report signatures of episodic positive selection (Alcaide and Edwards 2011; Grueber, et al. 2014; Outlaw, et al. 2019), suggesting diverse patterns of immune gene evolution. In *TLR* genes, a group of trans-membrane glycoproteins, sites of episodic positive selection are usually located in the extracellular ligand-binding domains of receptors that recognize highly conserved structural motifs expressed by microbial pathogens (i.e., pathogen-associated microbial patterns; PAMP; Alcaide and Edwards 2011; Brownlie and Allan 2011; Huang, et al. 2011). Different *TLR* identify specific types of PAMP and thus, functional changes at the gene level of different *TLR* genes are probably reflecting an adaptive response of the host immune system depending on the pathogen community (Khan, et al. 2019).

Despite recent advances (Minias, et al. 2017; O’Connor, et al. 2018), it remains generally unknown how pathogen-mediated selection affects immune gene evolution across species. Do signatures of positive selection mostly evolve convergent, or do they evolve divergent depending on pathogen exposure and life-history trade-offs? Recent work confirmed that pathogen exposure indeed is a key driver of pathogen-mediated selection of MHC genes (Minias, et al. 2017; O’Connor, et al. 2018). Ecological traits associated with increased pathogen exposure (migration, colonial breeding) positively affect selection strength on MHC genes (Minias, et al. 2017), but it remains unknown if this also is the case for *TLR* genes. In addition, life-history traits are likely to influence the strength of pathogen-mediated selection. This is because immunological responses are costly (Ricklefs and Wikelski 2002; Knowles, et al. 2009) and, therefore, hosts must trade-off their investment in immune functions and other life-history traits, including growth and reproduction. These trade-offs may limit resource allocation into immune functions. Thus, we predict that species with a lower investment into reproduction are more likely to have positively selected sites in *TLR* genes. To this end, we test for the role of ecology on molecular evolution of immune genes in a phylogenetic comparative framework, using data across almost all extant orders in birds.

Here we study the evolutionary patterns, and ecological and life-history drivers of avian *TLR* genes using a comparative approach. *TLRs* belong to an ancient gene family that is found across all animals and represent the first line of pathogen recognition and defense (Huang, et al. 2011). Although many studies have investigated evolutionary patterns of *TLR*s in some avian lineages (Alcaide and Edwards 2011; Brownlie and Allan 2011; Huang, et al. 2011; Grueber, et al. 2013; Grueber, et al. 2014; Velová, et al. 2018; Khan, et al. 2019), none of them have assessed their ecological drivers across all extant orders of birds. Compared to highly polymorphic MHC genes, which accordingly are difficult to genotype, *TLR* genes can be easily genotyped using designed universal primers (Alcaide and Edwards 2011; Grueber and Jamieson 2013). Moreover, whole genomes of many avian species are available, and birds are a highly diverse group with varying ecologies and life histories. We *de novo* sequenced *TLR* genes using a candidate gene approach and used published sequences to identify sites that are under positive selection, using selection analyses. Bases on these sites, we applied a comparative approach to assess the ecological and life-history traits associated with positive selection of avian *TLR* genes.

Based on previous works, we hypothesize that the evolution of *TLR* genes may be associated with ecological and life-history traits (Sheldon and Verhulst 1996; Minias, et al. 2017). We predict that species have more site under positive selection in their *TLR* genes when experiencing an increased pathogen exposure, which has been suggested to be the case in long-distance migrants (Altizer, et al. 2011), species with a larger breeding range (Altizer, et al. 2011), gregarious species (Altizer, et al. 2003), closed nesting species (Peralta-Sánchez, et al. 2012), species with a higher diet diversity (Benskin, et al. 2009), and polygamous species (Nunn 2002). Second, because immune responses are costly, they compete with other vital functions, including growth and reproduction (Lochmiller and Deerenberg 2000; Ricklefs and Wikelski 2002). Thus, we expect that altricial species, and species with a slow development, a long lifespan and extensive parental care (Lee, et al. 2008) have more sites of positive selection in their *TLR* genes compared to precocial species, and species with a fast development, a short lifespan and limited parental care.

## Results

### Evolutionary patterns of *TLR* genes in birds

Overall, we obtained DNA sequences of six *TLR* genes from 121 species from 58 families, covering 38 out of 40 extant avian orders (Table S2), selecting 1-3 species per order. Within species, the sequences in the paralogous *TLR1LA* and *TLR1LB* genes were more similar than those of other *TLR* genes (Table 1, Table S4-S5), supporting the previously shown occurrence of gene conversion in the extracellular domains of *TLR1LA* and *TLR1LB* (Alcaide and Edwards 2011). Such non-reciprocal transfer of gene sequences reflects unequal crossing-over between homologous sequences (Arnheim, et al. 1980), and is also found in immunoglobulin and major histocompatibility complex (*MHC*) gene families (McCormack, et al. 1991; Chaves, et al. 2010), which might be a way to increase the functional diversity of *TLR1* subfamily while minimizing genome size (Ohta 2010).

**Table 1.** Codons of Toll-like receptors genes (*TLR*) under positive or positive selection (ω > 1) at significance level *p* = 0.01, using three different methods: SLAC (Single Likelihood Ancestor Counting; Kosakovsky Pond and Frost 2005a), FEL (Fixed Effects Likelihood; Kosakovsky Pond and Frost 2005b), and MEME (Mixed Effects Model of Episodic Diversity Selection; Murrell, et al. 2012). The reported ω value were based on SLAC, representing overall mean ω value.

| Gene | Length (bp) | $\omega$ | SLAC | FEL | MEME |
| --- | --- | --- | --- | --- | --- |
| TLR1LA | 1176 | 0.389 | 37, 345 | 163, 167,<br>208, 214,<br>239 345 | 3, 37, 56, 89, 92, 232,<br>239, 242, 314, 338,<br>345, 356, 360, 365 |
| TLR1LB | 942 | 0.409 | 33, 81, 279 | 33, 81, 279 | 13, 21, 33, 40, 81, 166,<br>173, 272, 279, 308,<br>309, 311, 312, 313 |
| TLR3 | 906 | 0.327 | 212 | 217 | 84, 217, 302 |
| TLR4 | 867 | 0.492 | 72, 72, 101,<br>181, 200,<br>204, 205 | 72, 73, 127,<br>154, 172,<br>181, 200,<br>205 | 2, 3, 4, 72, 73, 127,<br>154, 172, 181, 200,<br>204, 205, 237, 246,<br>247, 286, 288 |
| TLR5 | 1254 | 0.452 | 95, 238 | 33, 95, 297,<br>343 | 5, 22, 33, 60, 95, 127,<br>131, 138, 220, 343,<br>415 |
| TLR7 | 1239 | 0.361 | 161, 190,<br>197, 344,<br>372 | 161, 190,<br>344, 372 | 24, 172, 190, 239,<br>280, 316, 344, 372,<br>411, 413 |

The phylogenetic trees of the six *TLR* genes (Figures 1A-F) recapitulated the phylogenetic relationships of major avian lineages, including the splits between Palaeognathae (ostrich and kiwi), Galloanseres (landfowl and waterfowl) and Neoaves (other orders). Within these major lineages, however, our analyses showed a high degree of incongruence between the *TLR* gene trees and a recent multi-locus avian phylogeny based on neutral genetic markers (Prum, et al. 2015), which may reflect stochastic lineage sorting during recent phases of fast radiation (Degnan and Rosenberg 2006; Suh, et al. 2015), or convergence evolution at different *TLR* loci (Králová, et al. 2018). Thus, ecological and life-history factors could play a role to mediate positive selection on *TLR* genes, which we explore in a second step with phylogenetic comparative analyses.

**Figure 1.**
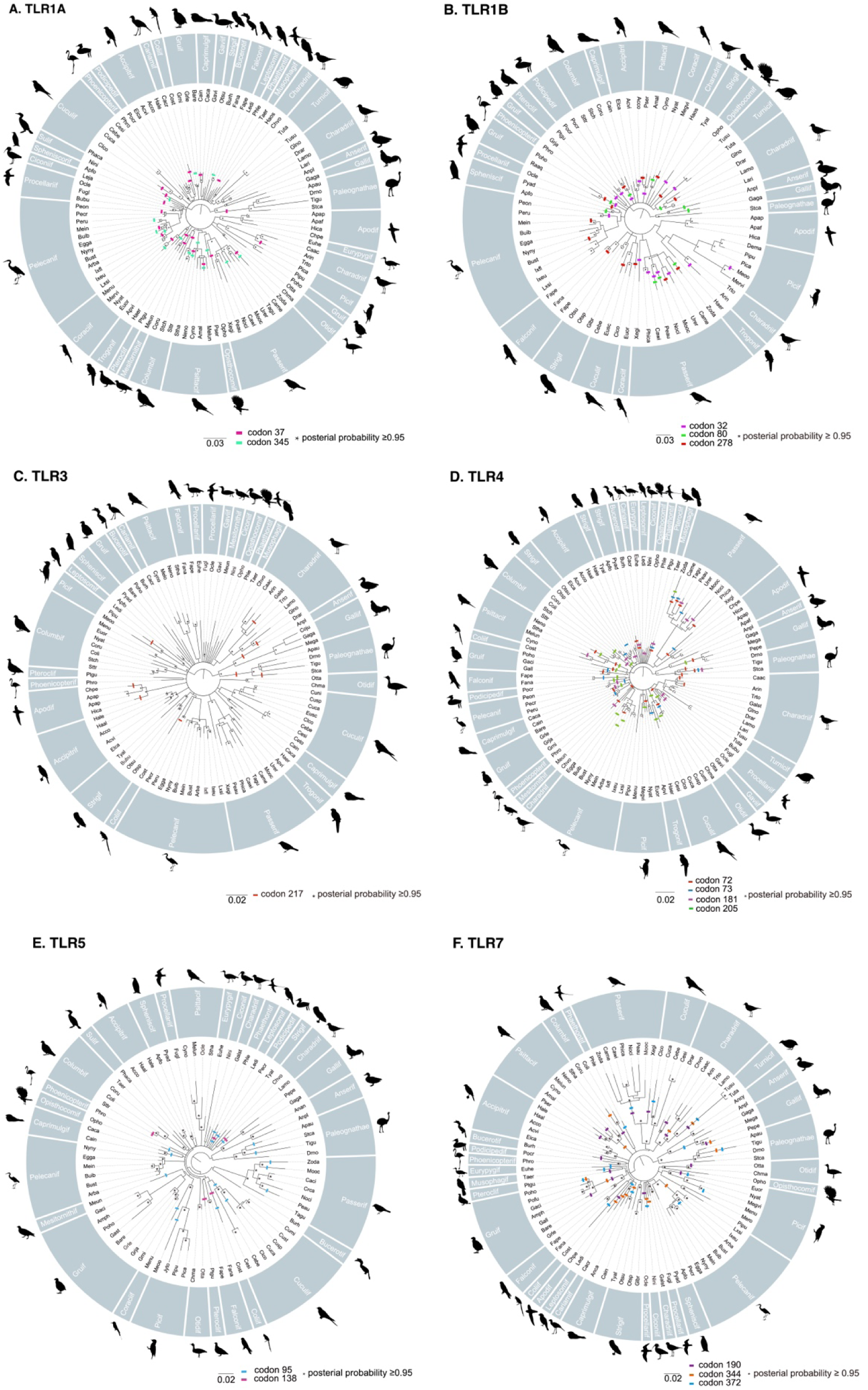
Phylogenetic reconstruction of the avian Toll-like receptors based on Bayesian inference. Codons with different color were the positive selection sites (ω > 1) detected by the software Datamonkey (Delport, et al. 2010) and clades with posterior probability are marked with an asterisk.

We established the selection patterns of *TLR* genes using three statistical approaches with different sensitivity and level of conservatism: SLAC (Single Likelihood Ancestor Counting; (Kosakovsky Pond and Frost 2005a), FEL (Fixed Effects Likelihood; Kosakovsky Pond and Frost 2005b) and MEME (Mixed Effects Model of Episodic Diversity Selection; Murrell, et al. 2012); see supplemental methods for details; Table S5). We found that the strength of selection ω (the rate of nonsynonymous substitutions (dN) divided by the rate of synonymous substitutions (dS), see methods) ranged between 0.33 and 0.49 for all *TLR* genes (Table 1; Figure 2). Accordingly, these genes are generally under purifying selection (Table 1). The number of sites under positive selection (ω > 1) differed depending on the used method, and were mostly located in the ligand-binding region of the ectodomain of *TLR* genes, indicating functional significance. The most conservative method (SLAC) identified 2-7 positively selected sites per *TLR* gene, which account for 0.16%∼0.81% of sequenced nucleotide sites, while a less conservative method (MEME) identified 3-17 positively selected sites per *TLR* gene (Table 1).

**Figure 2.**
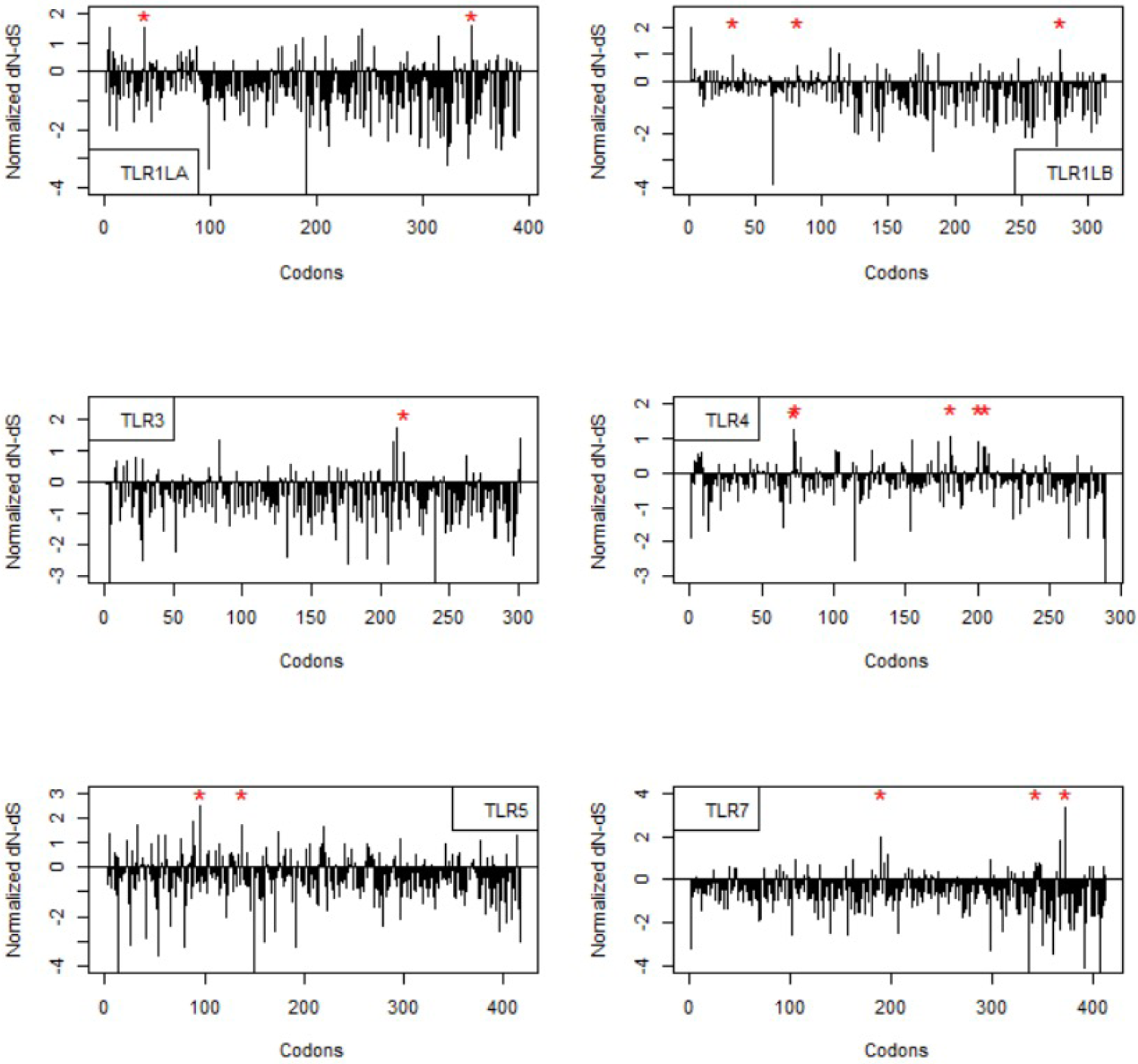
Positive selection sites in Toll-like receptors in 121 bird species. Site-wise rates of nonsynonymous and synonymous substitutions (dN/dS = ω) were calculated using Single Likelihood Ancestor Counting (Kosakovsky Pond and Frost 2005a). Sites under positive selection (ω > 1) are highlighted with a red asterisk.

### Ecological drivers of *TLR* gene evolution in birds

We calculated ω for each *TLR* gene using ancestral state reconstructions generated in PAML (Yang 2007). *TLR1LB* was excluded from these analyses given the high similarity between *TLR1LA* and *TLR1LB.* Phylogenetic comparative analyses (Figure 3, Table S7) showed that altricial species had higher values of ω than precocial species at *TLR1A* (Figure 4A), and that diet specialists had higher values of ω than food generalists at *TLR1A* (Figure 4B). Moreover, larger species had higher values of ω than smaller species at both *TLR4* (Figure 4D) and *TLR7* (Figure 4F). Family living species with prolonged parent-offspring associations had higher values of ω than non-family living species at *TLR4* (Figure 4C). Finally, long-distance migrants had higher values of ω than sedentary or partial migrants at *TLR5* (Figure 4E). Our analyses did not find an association between the value of ω and sexual dimorphism or breeding system (Figure 3).

**Figure 3.**
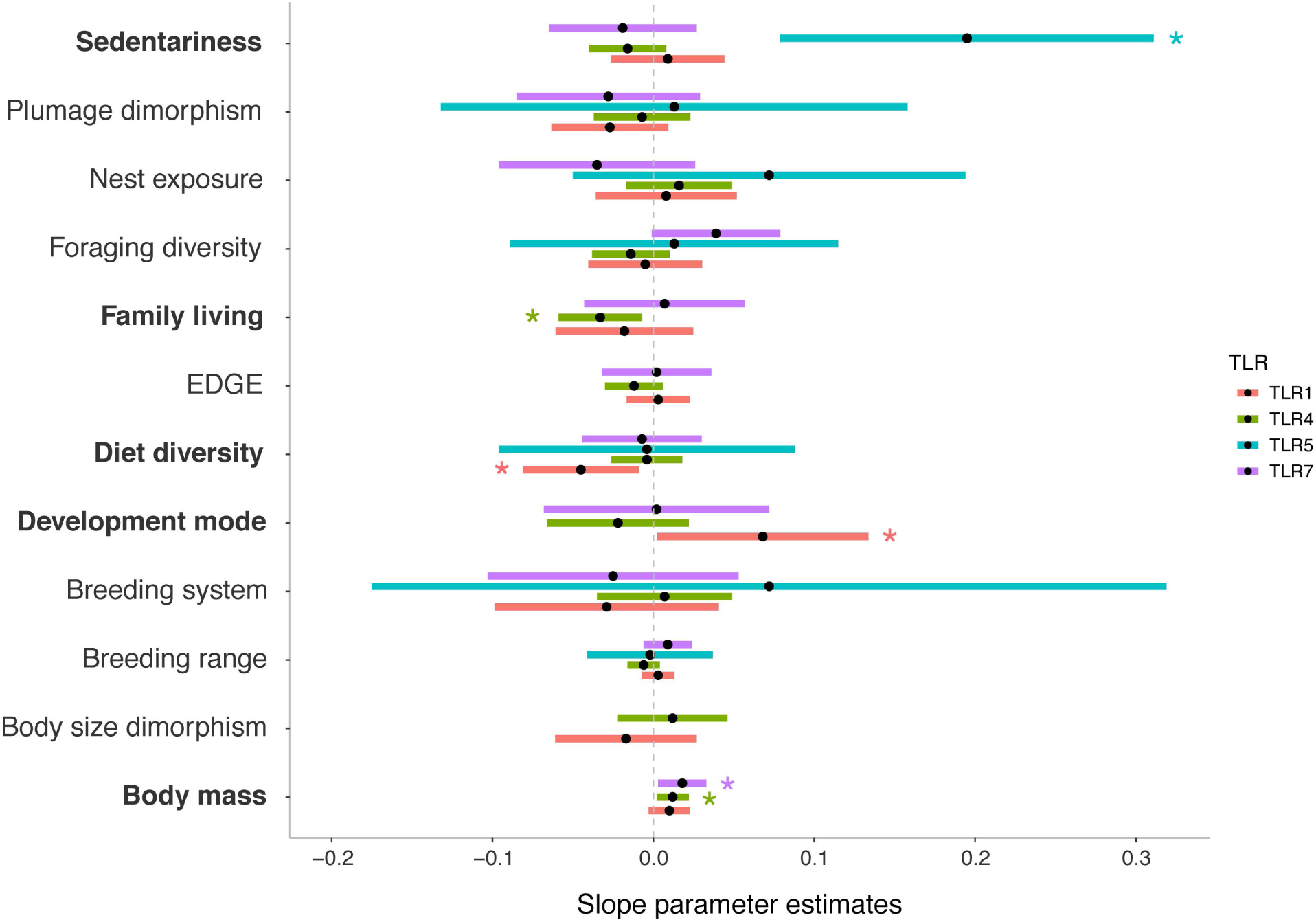
The posterior distributions (mean and 95% credible intervals) for the slope parameters for the averaged best-fitting models of the evolution of ω of four Toll-like receptor (*TLR*) genes. Significant parameters are marked with asterisk, and these parameters are highlighted in bold. *TLR1B* is not included in these analyses due to the high similarity between *TLR1LA* and *TLR1LB. TLR4* is not included in the figure as none of the factors is significant. Detailed information is found in Table S7.

**Figure 4.**
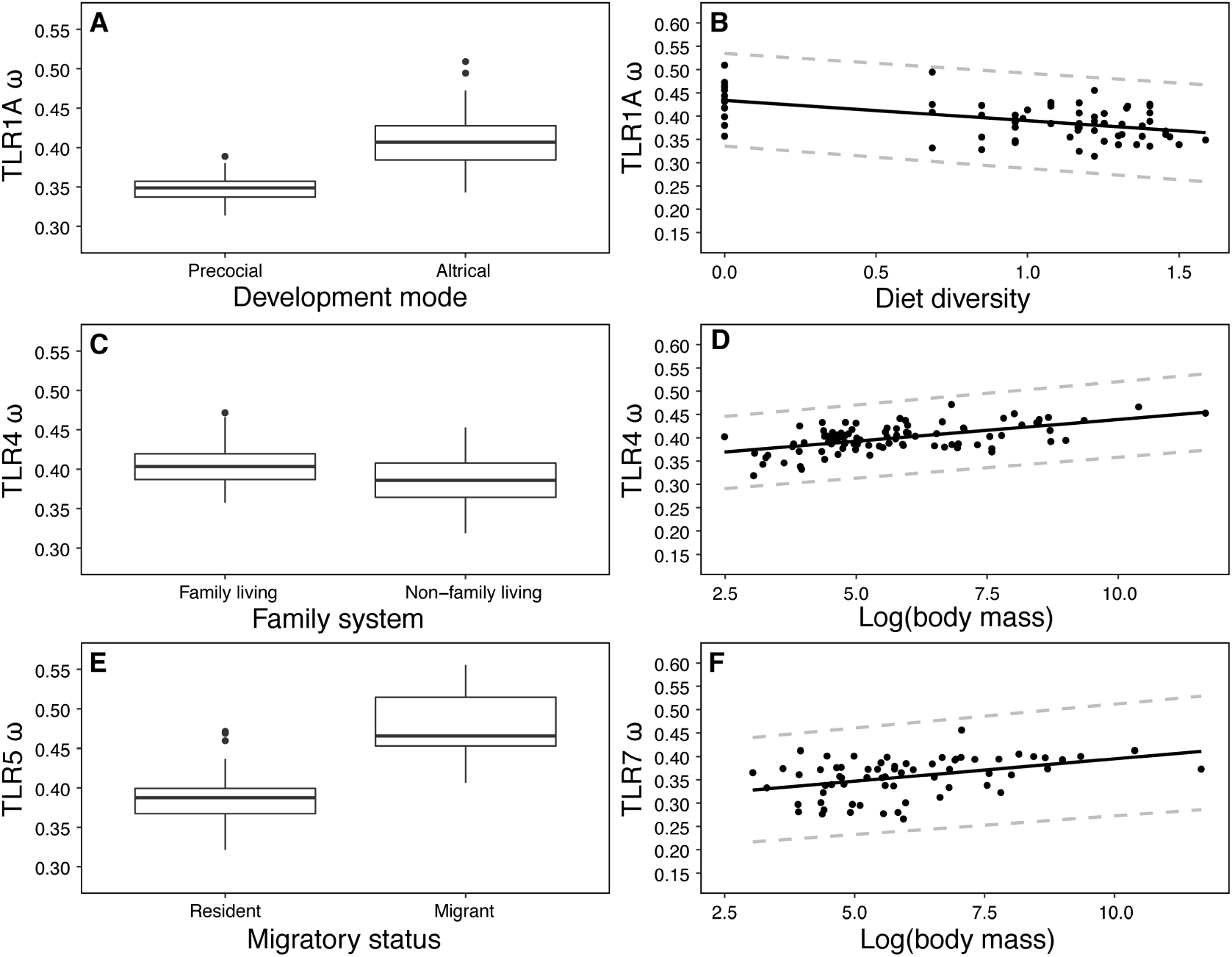
Significant ecological and life-history correlates are associated with positive selection (ω > 1) of the different Toll-like receptors in birds. Plots based on model predicted values based on phylogenetic mixed models. See also Table S7.

## Discussion

Our analyses of a phyla-wide data set of six avian *TLR* genes identified few positive selection sites in all *TLR* genes, supporting findings from single-species studies (Grueber, et al. 2014). The comparative analyses showed that positive selection in avian *TLR* genes reflects an increased exposure to pathogens and a life history involving extensive parental care. These findings complement recent studies, showing that MHC gene evolution is positively associated with migration and colonial breeding in birds (Minias, et al. 2017; Králová, et al. 2018; O’Connor, et al. 2018).

Most studies on immune gene evolution focused on single species and assessed the evolutionary and functional perspectives, while recently global analyses of micro- and macro-evolutionary patterns of immune genes emerged (Grueber, et al. 2014; Minias, et al. 2017; Velová, et al. 2018; Shultz and Sackton 2019). Our dataset of avian *TLRs* improves the resolution of discerning selection patterns and shows evidence of purifying selection. The few sites under episodic positive selection are located in the part of the of six *TLR* genes that encode the ectodomains. These results are consistent with previous studies in vertebrates, including primates (Wlasiuk and Nachman 2010), other mammals (Areal, et al. 2011), and several birds (Alcaide and Edwards 2011; Grueber, et al. 2014), reflecting evolutionary constraints to recognize PAMP (Roach, et al. 2005). The detected positive selected sites might be an adaptive response to pathogen-mediated selection to species-specific response to target PAMP of pathogens. Indeed, a recent study revealed that positively selected immune-related genes show up-regulated level of expression in response to pathogen challenges (Shultz and Sackton 2019).

Our results of the phylogenetic comparative analyses go beyond previous work and reveal associations between strength of selection and drivers from ecological and life-history factors in *TLR* genes. Consistent with previous studies (Nunn, et al. 2000; Nunn 2002), we showed that an increased risk to pathogen exposure can be associated with the evolution of a stronger immune response (Figure 3). However, our results further highlight that different *TLR* genes have specific ecological drivers related to differences in pathogen exposure. In particular, each *TLR* recognizes 1-2 pathogen categories that are associated with specific ecological and life-history factors (Nunn, et al. 2000; Nunn 2002), leading to different macroevolutionary dynamics of each *TLR* gene. For example, in *TLR5*, which is responsible for recognizing ligands from bacteria with flagella (Brownlie and Allan 2011), positive selection is favored by long-distance migration (Figure 4E). Long-distance migrants are exposed to a diverse spectrum of pathogens, including highly infectious bacteria (Slifko, et al. 2000; Webster, et al. 2005), increasing the need for efficient defenses against bacterial pathogens. This result is consistent with an avian MHC study (Minias, et al. 2017), and combined, these results suggest that pathogen exposure drives the evolution of both the innate and adaptive immune system response in birds. Furthermore, our results suggest that diet specialist have higher rates of positive selection than diet generalists in *TLR1A*. In birds, diet specialists have a lower physiological tolerance (Bonier, et al. 2007) and thus, could be more sensitive to immune challenges. This pattern may also explain why diet generalist are more likely to go extinct over evolutionary times compared to diet specialists (Burin, et al. 2016).

The expression of immune functions is energetically demanding, and likely to impose trade-offs between the investment in immune functions and other vital functions, including growth and reproduction. Our results suggest that altricial species (where young are provisioned and brooded by the parents after hatching) experience a higher rate of positive selection than precocial species (where young can feed, move and thermoregulation after hatching) at *TLR4* which identifies ligands from Gram-negative bacteria (including *E. coli*; Brownlie and Allan 2011). While precocial young are exposed to pathogens at an earlier stage of their life than altricial species, prolonged parental investment into young (Griesser, et al. 2017) seems to allow developing young to invest more energy into immune functions, and other costly traits, including larger brains (Isler and van Schaik 2012).

It has been suggested that sociality comes at the price of an increased risk of disease transmission (Altizer, et al. 2003). Individuals of species that form ephemeral flocks have increased interactions with conspecific and heterospecific individuals, exposing them to more diverse pathogens than species that live in stable family groups with prolonged parent-offspring associations (Griesser, et al. 2017). However, we found that family living species had higher positive selection at *TLR4* than non-family living species. Family living provides offspring with a safe haven allowing for protected acquisition of life skills (Griesser and Suzuki 2017), and social support from other group members (Almberg, et al. 2015). Consequently, offspring in family living species have a predictable access to resources (Covas and Griesser 2007), allowing juveniles to allocate more energy into the development the immune defense (as shown here). Thus, extended parental care may mitigate the social cost of directly transmitted infection (Altizer, et al. 2003), and this finding demonstrates a so far overlooked life-history trade-off between immune system evolution and energy available for development.

Finally, our results show that a heavier body weight is associated with stronger positive selection at *TLR1A* and *TLR7*. These receptors recognize ligands originated from mycobacteria and virus (including avian flu; Brownlie and Allan 2011) that affect populations of captive and wild birds (Tell, et al. 2001; Webster, et al. 2005). Given that heavier bird species are usually more long-lived (Valcu, et al. 2014), they have an increased risk of pathogen exposure, requiring a more efficient immune system. Moreover, heavier species have prolonged developmental periods, which may allow them to allocate more energy into the development of the immune system, as shown in several species of domesticated animals (Lochmiller and Deerenberg 2000), further supporting the development trade-off hypothesis. Thus, our results are consistent with a recent study showing that heavier species tend to have elevated rates molecular evolution (Iglesias-Carrasco, et al. 2019).

To conclude, our study reveals that evolutionary patterns of avian *TLR* genes include species-specific episodic positive selection leading to sequence changes at pathogen recognition sites, and pervasive purifying selection. Phylogenetic comparative analyses show that selection on *TLR* genes are associated with intraspecific variation in the exposure to pathogens and extensive parental care, highlighting the critical role of developmental constraints on the evolution of immune functions. Specific *TLR* genes are associated with identifying distinct pathogens, and diverse evolutionary history, including gene gain and loss (Velová, et al. 2018; Khan, et al. 2019), and different ecological and life-history drivers. These patterns of diversifying selection in avian *TLR* genes are associated with the rapid radiation of birds in diverse ecological niches and environments with different pathogen communities. Clearly, experimental work is needed to investigate the relationship between positive selected sites in *TLR* genes and individual fitness (Acevedo-Whitehouse, et al. 2005). Also population genetics studies will help to disentangle the relative role of nature selection and other evolutionary forces including genetic drift on *TLR* evolution (Grueber, et al. 2013). Overall our findings connect molecular evolution with macro-evolutionary processes, demonstrating the importance of involving hosts’ traits in understanding immune system evolution in birds.

## Materials and Methods

### Sample collection and DNA sequencing

We included 121 species from 38 bird orders in our study. We collected tissues or blood and isolated genomic DNA, tested the DNA quality, and thereafter amplified the target *TLR* genes in the same laboratory. For the DNA amplification and sequencing, we used published and specifically designed primers (Table S1). PCR products with a single band were sequenced in both directions using an ABI 3730XL Genetic Analyzer (BGI-Shenzhen). We applied candidate gene sequencing for the six *TLR* genes (N=263; see Table S2 for details), and combined these data with published DNA sequences (Alcaide and Edwards 2011; Huang, et al. 2011) from NCBI (N=530; http://www.ncbi.nlm.nih.gov).

We obtained *TLR* sequences from 121 species from 58 families, covering 38 out of 40 extant avian orders (Table S2), selecting 1-3 species per order. The sequence lengths varied between 867 base pairs (bp) in *TLR4* and 1254 bp in *TLR5* (Table S3). We included species with sequences from at least five *TLR* genes into the statistical analyses, yielding in about 60 species per gene.

### Phylogenetic analysis

Sequences were ensured, edited and aligned, visually checked and manually revised. We computed the best evolutionary model of nucleotide substitution of DNA sequence alignments and used Bayesian inference to reconstruct phylogenetic relationships with the associated nucleotide evolution model at each locus (Table S3).

Their phylogenetic relationships were reconstructed in the software MrBayes (Ronquist, et al. 2012) with parameters set as 15,000,000 generations using a general time reversible evolutionary model with gamma-distributed mutation rate variation across sites and a proportion of variable sites and a burn-in phase of 25%. The *TLR* genes of *Anas platyrhynchos* were used as outgroup. Gene conversion analysis was conducted in *TLR1LA* and *TLR1LB* using GARD (Genetic Algorithms for Recombination Detection; Pond, et al. 2006) and phylogenetic relationship was reconstructed with the recombination fragments.

### Tests of selection

To quantify positive selection strength, we estimated ω ((dN) / (dS)) in the package HyPhy (Pond and Muse 2005), using the Datamonkey web interface (http://www.datamonkey.org; Delport, et al. 2010). Values of ω > 1 suggest positive selection, while values of ω < 1 suggest purifying selection. We calculated ω values with three different approaches that differ in the underlying assumptions, using SLAC (Single Likelihood Ancestor Counting; Kosakovsky Pond and Frost 2005a), FEL (Fixed Effects Likelihood; Kosakovsky Pond and Frost 2005b), and MEME (Mixed Effects Model of Episodic Diversity Selection; Murrell, et al. 2012). SLAC is the most conservative method, which counts the number of nonsynonymous and synonymous substitutions along the phylogeny, by reconstructing the ancestral sequences. SLAC does not make any assumptions regarding the distribution of rates across sites and thus, underestimates the positive detection rate (Kosakovsky Pond and Frost 2005a). FEL fits substitution rates on a site-by-site basis, and make no assumption regarding the distribution of rates across sites (Kosakovsky Pond and Frost 2005b). Finally, MEME can detect instances of both episodic and pervasive positive selection at individual sites (Murrell, et al. 2012). We applied a conservative level of statistical significance at *p*=0.01 for these analyses.

### Phylogenetic comparative analysis

We used a maximum likelihood based method (Revell and Freckleton 2013) to estimate the ancestral state of ω of *TLR* distributions at each node using the platform Datamonkey (Delport, et al. 2010). Based on the ancestral state of ω, we estimated ω of every species using the default method implemented in PAML (YN00; Yang 2007), and used these values in our comparative analyses as response variable. Given the close phylogenetic link between *TLR1LA* and *TLR1LB*, we only included *TLR1LA TLR3, TLR4, TLR5* and *TLR7* in our analyses.

We used a model selection approach to investigate the influence of 12 different predictors on ω. Based on previous studies (Minias, et al. 2017; O’Connor, et al. 2018), we predict that the experienced infection risk and developmental trade-offs may influence ω. To assess the infection risk hypothesis, we included sedentariness (resident or short-distance migrant vs long-distance migrant), nest exposure (closed nests vs open nests), foraging habitat (water vs terrestrial) and a measurement of the dietary diversity. This latter parameter was calculated using the Shannon-Wiener index over all food items, following Wilman et al. (2014). To test the life history trade-off hypothesis, we included development mode (precocial, including semi-precocial) vs altricial, including semi-altricial), family system (no family living vs family living; see Drobniak, et al. 2015), breeding system (biparental care vs uniparental care), body size dimorphism (monomorphism vs dimorphism), plumage color dimorphism (monomorphism vs dimorphism), body mass (in grams), breeding range cell account, and a score of evolutionary distinctiveness of the species (EDGE score; Jetz, et al. 2014) as explanatory variables. Ecological and life-history parameters were obtained from bird handbooks (Del Hoyo, et al. 2011) and published studies (Jetz, et al. 2014; Valcu, et al. 2014; Griesser, et al. 2017).

The variance inflation factor (VIF) of all continuous parameters was well below 3, which indicates an acceptable amount of covariance among parameters (Dormann, et al. 2013). We log transformed non-normally distributed parameters (body mass, breeding cell count) to achieve normal distribution. We downloaded 10,000 phylogenetic trees for each *TLR* dataset from *BirdTree* (http://birdtree.org; Jetz, et al. 2012), and generated a maximum clade credibility tree (Figure S1 A-E) using TreeAnnotator v1.8.2 in the package BEAST (Drummond and Rambaut 2007). Bayesian phylogenetic mixed models with Gaussian error structure were run in the package MCMCglmm (Hadfield 2010), using a parameter expanded prior (V=1, nu=1, alpha.mu=0, alpha.V=1,000; Hadfield 2014). The MCMC chains were run for 75,000 iterations with a burn-in phase of 7,500 iterations and samples drawn every 300 iterations. Visual inspection of the MCMC chains revealed a low degree of autocorrelation and model convergence (Hadfield 2010).

We performed a model selection procedure including all explanatory variable listed above to identify the models with the highest explanatory degree. We ranked the candidate models according to their DIC (Deviance Information Criterion; Spiegelhalter, et al. 2002) using the “dredge” function of package MuMIn (Bartoń 2012). We report parameter estimates based on averaging the best fitting models (Δ DIC < 2; Burnham and Anderson 2002).

## Acknowledgements

We thank Pinjia Que, Qin Huang, Jian Zhao, Beijing Zoo and Qiyun Mountain National Natural Reserve for their invaluable sampling assistance, Szymon Drobniak for statistical advice and Carel van Schaik and Tamás Székely for helpful comments. This work was funded by the National Natural Science Foundation of China (No. 31301875 to Y.L.), Special Program for Applied Research on Super Computation of the NSFC-Guangdong Joint Fund (the second phase) under Grant No. U1501501 (to Y.L), a Swiss National Research Foundation travel grant on behalf of Y.L (to M.G.), and a Swiss National Research Foundation grant (PPOOP3_150752 to M.G.). Partial bird sampling works were supported by the Chinese National Science & Technology Basic Work Program, ‘‘The Comprehensive Scientific Survey of Biodiversity from Luoxiao Range Region in China (2013FY111500)’’ to YL.

