## Supplementary Information for "Infection risk and extensive parental care govern the molecular evolution of *Toll-like* receptors in birds"

[8]Senior authors.

[*] Lead contact

**This file includes:**

Supplementary text

Figures S1

Tables S1 to S7

SI References

Supplementary text

### Sample collection and DNA extraction

We collected tissue or blood samples from collections in Bio-museum, Sun Yat-sen University, South China Institute of Endangered Animals in Guangzhou, Zoological Institute of China Academy of Sciences (CAS) in Beijing, and Kunming institute of Zoology of CAS and Beijing Normal University Beijing. Besides, we downloaded 283 TLRs sequences in NCBI (National Center for Biotechnology Information, http://www.ncbi.nlm.nih.gov) database (Table S1). Totally, our study sample set covered 121 species of 58 families that belong to 38 orders of the Neognathae. We used the TIANamp Genomic DNA Kit to isolate genomic DNA (TIANGEN, Beijing, China) and tested DNA quality by Nanodrop 2000 Spectrophotometer (Nanodrop Technologies, Beijing, China).

### Gene amplification and sequencing

We referenced the published primers in Molecular Evolution of the Toll-like Receptor Multigene Family in Birds (Alcaide and Edwards 2011) and designed primers by Oligo v7.6.0 (Molecular Biology Insights, Inc., Cascade, Co) using *Gallus gallus* TLRs genes as reference. Primer properties (I.e. Tm, GC content) were investigated in software Oligo7. The list of primers used in this study was detailed as follows (Table S2).

TLRs gene amplification was performed in a 30-μL reaction volume containing approximately 20-100 ng genome DNA, 1 unit of Recombinant Taq DNA polymerase (TAKARA BIOTECHNOLOGY CO., DALIAN, China), 10×PCR buffer 3 μL, 50mM dNTP, 10 pmol of each primer. We carried out polymerase chain reaction (PCR) with the following temperature profiles: initial denaturation for 3 min at 94°C, followed by 35 cycles of 30s at 94°C, annealing for 30s at variable temperatures (Table S2), 80s at 72°C, with a final 10min elongation step at 72°C. PCR products were visualized at 1% agarose gels stained with SYBR Gold (Invitrogen, Shanghai, China) and sequenced in both directions on ABI 3730 XL Genetic Analyzer (service provided by Majorbio at Guangzhou, China).

**Statistical analysis**

Sequences were then imported into NCBI BLAST (Basic Local Search Tool, http://blast.ncbi.nlm.nih.gov/Blast) to test their homology with published avian *TLR* sequences. We further checked and did not found stop codons or frame shift mutations indicating genuine Toll-like receptor genes. DNA sequences were edited and aligned by DNASTAR Lasergene v7 (Burland 2000) and ClustalW v2.1 (MA, et al. 2007), checked by eye and manually revised if necessary by MEGA v6.06 (Tamura, et al. 2013). We introduced IUPAC (International Union of Pure and Applied Chemistry) degenerate nucleotides for those nucleotide positions found to be heterozygous in chromatograms. The alignments of nucleotides and amino acids for the analysis for gene conversion and phylogenetic analysis were using ClustalW and MEGA. jModelTest v.2.1.6 (Darriba, et al. 2012) and MrBayes v.3.2 (Ronquist, et al. 2012) were used to test the best evolutionary model of nucleotide substitution of DNA sequences alignments and construct the Bayesian inference of phylogenetic relationships respectively. The best model and the generations ran by MrBayes were listed in Table S3.

To conduct gene conversion analysis, we selected species that were genotyped at both *TLR1LA* and *TLR1LB* genes. Recombination analysis was performed using GENCONV v.1.81 (Sawyer 1989) with default settings. The more conservative and accurate p-values were used with 10,000 pseudo-replicates permutations.

**Ecological traits collection**

To test whether there is any relationship between microevolution and macroevolution, we used ω (the rate of nonsynonymous substitutions (dN) divided by the rate of synonymous substitutions (dS) of each *TLR* as response variable and nine environment variables (chicken development mode, family system, breeding system, migration, sexual selection, foraging mode, body weight, BRCC (Breeding Range Cell Account) (Jetz, et al. 2014), EDGE (Evolutionary Distinct and Globally Endangered score) (Jetz, et al. 2014), range) as explanatory variables. Correlation test among continuous variables (BRCC, EDGE and body weight) were conducted and screened the autocorrelation variables. Besides, these continuous variables were tested whether they were normal distribution. In this study, body weight and ω of *TLR3* were log-transformed to be normal.

**Gene conversion detection**

We found that the two paralogues genes seemed to show sequence similarity. We chose species that both sequenced *TLR1LA* and *TLR1LB* to detect whether there existed gene conversion in birds. We found strong evidence of gene conversion that a highly significance value was derived. It is apparently that the paralogues genes were clustered by species rather than by locus, indicating that the paralogues genes were more resemble with each other.

The distinction of amino acid sequences remains changeless throughout evolution or became identical through convergent evolution and sequences turn out to be identical by gene conversion is that the latter were encoded by the same codons. To discriminate whether the cause of invariable sequences, we analyzed the codon usage in regions according to the outcome of GENCONV. The averaged codon usage in the conserved codons in the possible gene conversion region was up to 99% or 100%. Taking together, we found that gene conversion was the most possible procession that give rise to the changeless codons between the two paralogues.

**Figure S1 a-e** show the distribution of Neoaves, (a)TLR1LA, (b)TLR3, (c)TLR4, (d)TLR5, (e)TLR7 along phylogenies of studied avian species.

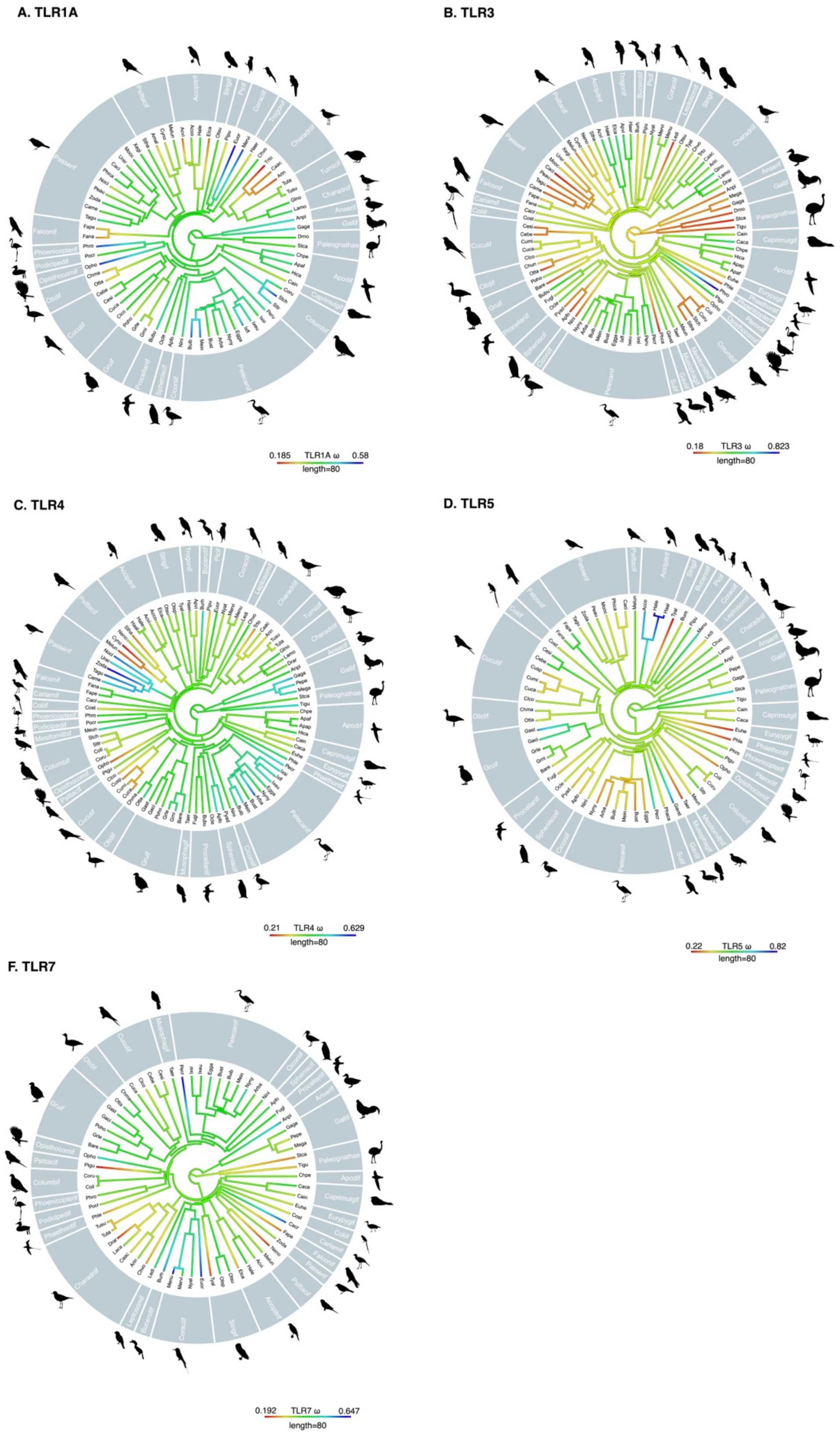

**Table S1** Primers designed in this study and associated annealing temperature (Ta) used to amplify *TLR1LA*, *TLR1LB* and *TLR3*. Lengths shown are total amplicon length, including primers and are approximate. Related to Key Resources Table in the main text.

| Genes | Primers | Sequences (5’-3’) | Ta (℃) | Length (bp) |
|---|---|---|---|---|
| *TLR1LA* | 865F | CTTATATGATGGAATGAGCACT | 56.5 | 1494 |
| | 2338R | TTGTAGCTCTTCTCAATGCAG | | |
| | 942F | CCTCCCCTTCTTTAGAGCTT | 57 | 1478 |
| | 2400R | GTACAGCTCGTAGTGACACC | | |
| | 979F | AACAGATCTAACGCTTGACAC | 57 | 1445 |
| | 2403R | CAAAGTACAGCTCGTAGTGAC | | |
| | 803F | GCTTTAAATACACGGAGCCT | 56.5 | 1339 |
| | 2123R | TCCTCTTCGTCTGCGTCCA | | |
| *TLR1LB* | 1090F | ATGAATATTGCAGCCTTGACA | 57.5 | 1163 |
| | 2234R | TACAGCTCGTAGTGACACC | | |
| | 1145F | CTTCATCTGACAGTCCCCTT | 58 | 1109 |
| | 2234R | GTACAGCTCGTAGTGACACC | | |
| | 905F | TACTTCAGGTTGTATGGCACT | 57 | 1071 |
| | 1957R | TCCTCTTCGTCTGCGTCCA | | |
| *TLR3* | TLR3-F-2 | GAATTATCAAACACAGCGAT | 54 | / |

**Table S2** NCBI accession numbers and sequenced genes of Toll-like receptors included in this study.

"/" represent unsubmitted sequences generated from this study. GenBank accession numbers will be available upon acceptance of the manuscript.

| **Species** | Abbreviation | Order | *TLR1LA* | *TLR1LB* | *TLR3* | *TLR4* | *TLR5* | *TLR7* |
|---|---|---|---|---|---|---|---|---|
| ***Accipiter cooperii*** | Acco | Accipitriformes | GU904995 | GU904946 | HQ267385 | GU904970 | GU904974 | GU904982 |
| ***Accipiter virgatus*** | Acvi | Accipitriformes | | | | | | |
| ***Elanus caeruleus*** | Elca | Accipitriformes | | | | | | |
| ***Haliaeetus albicilla*** | Haal | Accipitriformes | | | XM_009915940.1 | XM_009915776.1 | XM_009928565.1 | XM_009913820.1 |
| ***Haliaeetus leucocephalus*** | Hale | Accipitriformes | XM_010574438.1 | / | XM_010570028.1 | XM_010571006.1 | XM_010577687.1 | XM_010581565.1 |
| ***Anser anser*** | Anan | Anseriformes | / | / | / | / | JX096947 | / |
| ***Anser cygnoides*** | Ancy | Anseriformes | / | / | / | / | / | JQ910168 |
| ***Anas platyrgynchos*** | Anpl | Anseriformes | FJ477859.1 | FJ477859.1 | JN573268.1 | JQ839148 | JN573267.1 | JQ687402.1 |
| ***Apus affinis*** | Apaf | Apodiformes | | | | | / | / |
| ***Apus apus*** | Apap | Apodiformes | | | | | / | / |
| ***Chaetura pelagica*** | Chpe | Apodiformes | XM_010006120.1 | | XM_010002467.1 | XM_010007952.1 | / | XM_010008803.1 |
| ***Hirundapus caudacutus*** | Hica | Apodiformes | | | | | / | / |
| ***Buceros rhinoceros*** | Burh | Bucerotiformes | XM_010135841.1 | | XM_010135016.1 | XM_010137451.1 | XM_010140290.1 | XM_010140535.1 |
| ***Caprimulgus carolinensis*** | Caca | Caprimulgiformes | XM_010167982.1 | | XM_010170240.1 | XM_010165071.1 | XM_010164361.1 | XM_010175260.1 |
| ***Caprimulgus indicus*** | Cain | Caprimulgiformes | | | | | | |
| ***Cariama cristata*** | Cacr | Cariamiformes | XM_009699424.1 | | XM_009700362.1 | XM_009702452.1 | / | XM_009696068.1 |
| ***Dromaius novaehollandia*** | Drno | Casuariiformes | GU904989 | / | GU904961 | GU904966 | GU904972 | GU904977 |
| ***Arenaria interpres*** | Arin | Charadriiformes | | | | | / | |
| ***Calidris acuminata*** | Caac | Charadriiformes | | / | | | / | |
| ***Charadrius vociferus*** | Chvo | Charadriiformes | XM_009884994.1 | / | XM_009883901.1 | XM_009888044.1 | XM_009894296.1 | XM_009895207.1 |
| ***Dromas ardeola*** | Drar | Charadriiformes | | | | | / | |
| ***Gallinago stenura*** | Gast | Charadriiformes | / | / | | | / | / |
| ***Glareola nordmanni*** | Glno | Charadriiformes | | | | | / | / |
| ***Haematopus ostralegus*** | Haos | Charadriiformes | | | / | / | / | / |
| ***Larus mongolicus*** | Lamo | Charadriiformes | | | | | | |

| | | | | | | | | |
|---|---|---|---|---|---|---|---|---|
| ***Larus ridibundus*** | Lari | Charadriiformes | | | / | | / | / |
| ***Tringa totanus*** | Trto | Charadriiformes | | | | | / | |
| ***Colius striatus*** | Cost | Coliiformes | XM_010205936.1 | / | XM_010203505.1 | XM_010195927.1 | XM_010204457.1 | XM_010209014.1 |
| ***Columba livia*** | Coli | Columbiformes | / | / | XM_005500210.1 | XM_005498384.1 | XM_005511337.1 | XM_005512700.1 |
| ***Columba rupestris*** | Coru | Columbiformes | | | | | | |
| ***Streptopelia chinensis*** | Stch | Columbiformes | | | | | / | / |
| ***Streptopelia tranquebarica*** | Sttr | Columbiformes | | | | | | / |
| ***Eurystomus orientalis*** | Euor | Coraciiformes | | | / | | / | |
| ***Merops nubicus*** | Menu | Coraciiformes | XM_008949981.1 | / | XM_008943939.1 | XM_008945045.1 | XM_008950209.1 | XM_008935272.1 |
| ***Merops viridis*** | Mevi | Coraciiformes | | | | | / | |
| ***Nyctyornis athertoni*** | Nyat | Coraciiformes | | | | | / | |
| ***Centropus bengalensis*** | Cebe | Cuculiformes | | | | / | | |
| ***Centropus sinensis*** | Cesi | Cuculiformes | | / | | | | |
| ***Clamator coromandus*** | Clco | Cuculiformes | | | | | | |
| ***Cuculus canorus*** | Cuca | Cuculiformes | XM_009566580.1 | / | XM_009565656.1 | XM_009569425.1 | XM_009569717.1 | XM_009555971.1 |
| ***Cuculus micropterus*** | Cumi | Cuculiformes | / | / | | | | / |
| ***Cuculus sparverioides*** | Cusp | Cuculiformes | / | / | | | | / |
| ***Centropus toulou*** | Ceto | Cuculiformes | / | / | | / | / | / |
| ***Eudynamys scolopacea*** | Eusc | Cuculiformes | / | | | / | / | / |
| ***Nipponia nippon*** | Nini | Ciconiiformes | XM_009461836.1 | / | XM_009474837.1 | XM_009475802.1 | XM_009472928.1 | XM_009476283.1 |
| ***Eurypyga helias*** | Euhe | Eurypygiformes | XM_010158154.1 | / | XM_010156014.1 | XM_010156250.1 | XM_010151494.1 | XM_010146638.1 |
| ***Falco naumanni*** | Fana | Falconiformes | GU904861 | GU904872 | GU904898 | GU904900 | GU904907 | GU904923 |
| ***Falco peregrinus*** | Fape | Falconiformes | XM_005243208.2 | XM_005244547.1 | XM_005243141.1 | XM_005231393.1 | XM_005241848.1 | XM_005229443.1 |
| ***Gallus gallus*** | Gaga | Galliformes | NM_001007488 | NM_001081709 | NM_001011691 | NM_001030693 | NM_001024586 | NM_001011688 |
| ***Meleagris gallopavo*** | Mega | Galliformes | / | / | XM_003205774.2 | XM_003211211.2 | / | XM_003203086 |
| ***Perdix perdix*** | Pepe | Galliformes | / | / | / | JQ713172 | JQ713180 | JQ713178 |
| ***Gavia stellata*** | Gavi | Gaviiformes | / | / | XM_009811989.1 | XM_009809127.1 | XM_009821515.1 | XM_009813692.1 |
| ***Balearica regulorum*** | Bare | Gruiformes | XM_010306228.1 | / | XM_010310965.1 | XM_010304091.1 | XM_010301313.1 | XM_010309901.1 |
| ***Grus japonensis*** | Grja | Gruiformes | / | | / | | | / |

| | | | | | | | | |
|---|---|---|---|---|---|---|---|---|
| ***Grus leucogeranus*** | Grle | Gruiformes | | / | / | | | |
| ***Grus nigricollis*** | Grni | Gruiformes | | / | / | | | |
| ***Porphyrio hochstetteri*** | Poho | Gruiformes | KF265263 | KF265273 | KF265288 | KF265299 | KF265312 | KF265323 |
| ***Rallus aquaticus*** | Raaq | Gruiformes | / | | / | / | / | / |
| ***Amaurornis phoenicurus*** | Amph | Gruiformes | / | / | / | / | | |
| ***Gallicrex cinerea*** | Gaci | Gruiformes | / | / | / | | | |
| ***Gallirallus striatus*** | Gall | Gruiformes | / | / | / | | | |
| ***Porzana fusca*** | Pofu | Gruiformes | / | / | / | / | / | |
| ***Leptosomus discolor*** | Ledi | Leptosomiformes | XM_009960968.1 | / | XM_009953213.1 | XM_009947763.1 | XM_009950947.1 | XM_009960331.1 |
| ***Mesitornis unicolor*** | Meun | Mesitornithiformes | XM_010178919.1 | / | XM_010181262.1 | XM_010193770.1 | XM_010194240.1 | / |
| ***Tauraco erythrolophus*** | Taer | Musophagiformes | XM_009981391.1 | / | XM_009986371.1 | XM_009978945.1 | XM_009978079.1 | XM_009991025.1 |
| ***Ophisthocomus hoazin*** | Opho | Opisthocomiformes | XM_009935562.1 | XM_009943145.1 | XM_009940192.1 | XM_009935409.1 | XM_009937341.1 | XM_009943495.1 |
| ***Chlamydotis macqueenii*** | Chma | Otidiformes | XM_010124043.1 | / | XM_010122732.1 | XM_010127205.1 | XM_010119531.1 | XM_010119851.1 |
| ***Otis tarda*** | Otta | Otidiformes | | / | | | | |
| ***Callaeas wilsoni*** | Cawi | Passeriformes | KF265256 | KF265266 | KF265283 | / | KF265304 | KF265316 |
| ***Carpodacus mexicanus*** | Came | Passeriformes | GU904709 | GU904771 | GU904804 | GU904813 | / | GU904828 |
| ***Philesturnus carunculatus*** | Phca | Passeriformes | KF265261 | KF265272 | | | | |
| ***Mohoua ochrocephala*** | Mooc | Passeriformes | KF265259 | KF265269 | KF265285 | KF265293 | KF265308 | KF265319 |
| ***Notiomystis cincta*** | Noci | Passeriformes | KF265260 | KF265271 | / | KF265295 | KF265310 | KF265320 |
| ***Petroica australis rakiura*** | Peau | Passeriformes | JX502626 | JX502629 | JX502638 | JX502640 | JX502646 | JX502658 |
| ***Taeniopygia guttata*** | Tagu | Passeriformes | NW_002198637: 1935740_1938669 | NW_002198637: 1925319-1926697 | NW_002198636: 2625536-2631659 | NM_001142454 | NW_002198506: 243510-246064 | NW_002197669: 14707528-14718808 |
| ***Urocissa erythrorhyncha*** | Urer | Passeriformes | | | | | / | / |
| ***Xenicus gilviventris*** | Xegi | Passeriformes | KF265265 | KF265274 | KF265290 | KF265301 | / | KF265325 |
| ***Zoothera dauma*** | Zoda | Passeriformes | | | / | | | |
| ***Ardeola bacchus*** | Arba | Pelecaniformes | | / | | | | |
| ***Bubulcus ibis*** | Buib | Pelecaniformes | | | | | | |
| ***Butorides striatus*** | Bust | Pelecaniformes | | | | | | |
| ***Egretta garzetta*** | Egga | Pelecaniformes | XM_009641277.1 | XM_009641275.1 | XM_009642418.1 | XM_009634876.1 | XM_009637106.1 | XM_009646337.1 |
| ***Ixobrychus eurhythmus*** | Ixeu | Pelecaniformes | | | | | / | |

| | | | | | | | | |
|---|---|---|---|---|---|---|---|---|
| ***Ixobrychus flavicollis*** | Ixfl | Pelecaniformes | | | | | / | / |
| ***Ixobrychus sinensis*** | Lxsi | Pelecaniformes | | | | | / | |
| ***Mesophoyx intermedia*** | Mein | Pelecaniformes | | | | | | |
| ***Nycticorax nycticorax*** | Nyny | Pelecaniformes | | | | | | |
| ***Pelecanus crispus*** | Pecr | Pelecaniformes | XM_009478351.1 | / | XM_009486220.1 | XM_009481688.1 | XM_009485400.1 | XM_009489888.1 |
| ***Pelecanus onocrotalus*** | Peon | Pelecaniformes | | | / | | / | / |
| ***Pelecanus rufescens*** | Peru | Pelecaniformes | | | | | / | / |
| ***Phaethon lepturus*** | Phle | Phaethontiformes | XM_010285878.1 | / | XM_010281742.1 | XM_010293516.1 | XM_010295212.1 | XM_010290259.1 |
| ***Phoenicopterus roseus*** | Phro | Phoenicopteriformes | | | | | | |
| ***Dendrocopos major*** | Dema | Piciformes | / | | / | / | / | / |
| ***Jynx torquilla*** | Jyto | Piciformes | / | / | / | / | | / |
| ***Megalaima oorti*** | Meoo | Piciformes | / | | | / | | / |
| ***Megalaima virens*** | Mevi | Piciformes | / | | / | / | / | |
| ***Picus canus*** | Pica | Piciformes | / | | / | / | | / |
| ***Picoides pubescens*** | Pipu | Piciformes | GU904994 | GU904949 | GU904964 | GU904971 | GU904976 | GU904983 |
| ***Podiceps cristatus*** | Pocr | Podicipediformes | | | / | | / | |
| ***Bulweria bulwerii*** | Bubu | Procellariiformes | | / | | | / | / |
| ***Fulmarus glacialis*** | Fugl | Procellariiformes | XM_009583076.1 | / | XM_009585392.1 | XM_009586474.1 | XM_009572390.1 | XM_009572361.1 |
| ***Oceabdroma leucorhoa*** | Ocle | Procellariiformes | GU904993 | GU904947 | GU904965.1 | GU904969 | GU904975 | GU904981 |
| ***Amazona albifrons*** | Amal | Psittaciformes | GU904992 | GU904948 | / | / | / | GU904980 |
| ***Cyanoramphus novaezelandiae*** | Cyno | Psittaciformes | KF265257 | KF265267 | KF265284 | KF265292 | KF265307 | KF265318 |
| ***Melopsittacus undulatus*** | Melo | Psittaciformes | XM_005149134.1 | XM_013128310.1 | XM_005149053.2 | XM_005145616.1 | XM_005151875.1 | XM_005151439.1 |
| ***Nestor notabilis*** | Neno | Psittaciformes | XM_010022813.1 | / | XM_010013832.1 | XM_010014853.1 | / | XM_010019975.1 |
| ***Psittacus erithacus*** | Pser | Psittaciformes | | | / | / | / | |
| ***Strigops habroptila*** | Stha | Psittaciformes | KF265264 | / | KF265289 | KF265300 | KF265313 | KF265324 |
| ***Pterocles gutturalis*** | Ptgu | Psittaciformes | XM_010077749.1 | XM_010086209.1 | XM_010074246.1 | XM_010073689.1 | XM_010074631.1 | XM_010076693.1 |
| ***Aptenodytes forsteri*** | Apfo | Sphenisciformes | XM_009280175.1 | XM_009280152.1 | XM_009277378.1 | XM_009282256.1 | XM_009275754.1 | XM_009278529.1 |

| | | | | | | | | |
|---|---|---|---|---|---|---|---|---|
| ***Pygoscelis adeliae*** | Pyad | Sphenisciformes | / | XM_009332470.1 | XM_009330657.1 | XM_009319316.1 | XM_009333665.1 | XM_009318873.1 |
| ***Glaucidium brodiei*** | Glbr | Strigiformes | / | | / | / | / | |
| ***Otus spilocephalus*** | Otsp | Strigiformes | / | | | | / | |
| ***Otus sunia*** | Otsu | Strigiformes | | | | | / | |
| ***Tyto alba*** | Tyal | Strigiformes | / | XM_009975024.1 | XM_009970053.1 | XM_009970420.1 | XM_009968647.1 | XM_009975060.1 |
| ***Struthio camelus*** | Stca | Struthioniformes | XM_009673363.1 | XM_009688066.1 | XM_009676700.1 | XM_009668178.1 | XM_009686697.1 | XM_009683661.1 |
| ***Phalacrocorax carbo*** | Phca | Suliformes | XM_009511286.1 | / | XM_009505917.1 | XM_009514274.1 | XM_009513901.1 | XM_009501983.1 |
| ***Tinamus guttatus*** | Tigu | Tinamiformes | XM_010219197.1 | / | XM_010219604.1 | XM_010219200.1 | XM_010217273.1 | XM_010228317.1 |
| ***Apaloderma vittatum*** | Apvi | Trogoniformes | XM_009877018.1 | / | XM_009871058.1 | XM_009867818.1 | / | / |
| ***Harpactes erythrocephalus*** | Haer | Trogoniformes | | | | | / | / |
| ***Turnix suscitator*** | Tusu | Turniciformes | | | / | | / | |
| ***Turnix tanki*** | Tuta | Turniciformes | | | / | | / | |

**Table S3** Sequence alignments, best evolutionary substitution model and generations ran in MrBayes.

| Gene | Length (bp) | No. of species | Model | Generations |
|---|---|---|---|---|
| *TLR1LA* | 1176 | 100 | SYM+I+G | 15 M |
| *TLR1LB* | 942 | 75 | GTR+G | 8 M |
| *TLR3* | 906 | 97 | GTR+I+G | 5 M |
| *TLR4* | 867 | 104 | SYM+I+G | 8 M |
| *TLR5* | 1254 | 77 | GTR+I+G | 8 M |
| *TLR7* | 1239 | 93 | GTR+I+G | 8 M |

**Table S4** Gene conversion analysis for *TLR1LA* and *TLR1LB* sequences for 69 birds. Abbreviation: species names, see Table S2; BC KA: Bonferroni-corrected KA (BLAST-like) P-values. KA P values are NOT corrected for multiple pairwise comparisons. BC P-values: KA P-values multiplied by 9453. Num Poly: number of polymorphic sites in the fragment. Num Dif: number of mismatches within the fragment. Tot Difs: total number of mismatches between two sequences.

| Species | BC KA p-value | Begin | End | Length | Num Poly | Num Dif | Tot Difs |
|---|---|---|---|---|---|---|---|
| Poho | <0.00001 | 288 | 912 | 625 | 374 | 0 | 139 |
| Mero | <0.00001 | 316 | 912 | 610 | 365 | 0 | 135 |
| Apap | <0.00001 | 286 | 912 | 627 | 376 | 0 | 130 |
| Arin | <0.00001 | 311 | 832 | 522 | 336 | 0 | 140 |
| Trto | <0.00001 | 386 | 912 | 527 | 315 | 0 | 146 |
| Egga | <0.00001 | 291 | 873 | 583 | 370 | 0 | 122 |
| Coru | <0.00001 | 291 | 912 | 622 | 371 | 0 | 120 |
| Nyny | <0.00001 | 324 | 912 | 589 | 353 | 0 | 120 |
| Apaf | <0.00001 | 385 | 873 | 489 | 315 | 0 | 120 |
| Apfo | <0.00001 | 387 | 912 | 526 | 314 | 0 | 127 |
| Pser | <0.00001 | 398 | 912 | 515 | 306 | 0 | 128 |
| Pser | 0.27 | 291 | 394 | 104 | 64 | 0 | 128 |
| Glno | <0.00001 | 399 | 797 | 399 | 256 | 0 | 130 |
| Glno | 0.16 | 291 | 397 | 107 | 65 | 0 | 130 |
| Haos | <0.00001 | 532 | 912 | 381 | 219 | 0 | 129 |
| Haos | 0.28 | 417 | 530 | 114 | 73 | 0 | 129 |
| Elca | <0.00001 | 283 | 607 | 325 | 203 | 0 | 129 |
| Stch | <0.00001 | 286 | 623 | 338 | 212 | 0 | 122 |
| Hica | <0.00001 | 577 | 912 | 336 | 191 | 0 | 134 |
| Hica | <0.00001 | 336 | 575 | 240 | 154 | 0 | 134 |
| Pocr | <0.00001 | 291 | 605 | 315 | 193 | 0 | 126 |
| Pocr | 0.0003 | 607 | 749 | 143 | 94 | 0 | 126 |
| Otsu | <0.00001 | 556 | 857 | 302 | 187 | 0 | 126 |
| Acvi | <0.00001 | 461 | 740 | 280 | 180 | 0 | 129 |
| Acvi | <0.00001 | 283 | 459 | 177 | 110 | 0 | 129 |
| Sttr | <0.00001 | 585 | 841 | 257 | 167 | 0 | 126 |
| Sttr | 0.005 | 385 | 508 | 124 | 82 | 0 | 126 |
| Clco | <0.00001 | 639 | 912 | 274 | 153 | 0 | 136 |
| Opho | <0.00001 | 650 | 912 | 263 | 147 | 0 | 133 |
| Opho | <0.00001 | 426 | 617 | 192 | 118 | 0 | 133 |
| Anpl | <0.00001 | 470 | 658 | 189 | 123 | 0 | 143 |
| Anpl | 0.34 | 381 | 468 | 88 | 56 | 0 | 143 |
| Cain | 0.028 | 658 | 765 | 108 | 70 | 0 | 134 |
| Cain | 0.036 | 767 | 873 | 107 | 69 | 0 | 134 |

**Table S5** Codon usage for conserved amino acids between *TLR1LA* and *TLR1LB* in amino acid region under gene conversion. For all conserved amino acids, whether the codon used in TLR1LA or TLR1LB was identical or synonymous. The results were presented as identical/ (identical + synonymous). Species name abbreviations were referred Table S2.

| Species | Identical codons | Conserve codons | Codon usage (%) |
|---|---|---|---|
| Poho | 204 | 204 | 100 |
| Mero | 203 | 204 | 99 |
| Apap | 204 | 204 | 100 |
| Arin | 201 | 204 | 98 |
| Trto | 204 | 204 | 100 |
| Egga | 196 | 197 | 99 |
| Coru | 203 | 203 | 100 |
| Nyny | 202 | 202 | 100 |
| Apaf | 202 | 203 | 99 |
| Apfo | 201 | 202 | 99 |
| Pser | 202 | 203 | 99 |
| Glno | 201 | 201 | 100 |
| Fape | 200 | 202 | 99 |
| Haos | 198 | 201 | 98 |
| Elca | 202 | 203 | 99 |
| Stch | 202 | 203 | 99 |
| Hica | 201 | 202 | 99 |
| Pocr | 200 | 202 | 99 |
| Otsu | 198 | 200 | 99 |
| Acvi | 200 | 202 | 99 |
| Sttr | 196 | 199 | 98 |
| Clco | 199 | 202 | 98 |
| Opho | 199 | 201 | 99 |
| Anpl | 198 | 199 | 99 |
| Cain | 197 | 201 | 98 |

**Table S6** Dataset of Neoaves of TLRs and trait values used in comparative analysis.

The abbreviations are: CM = chick modus: A: precocial, B: altrical; FS = family system: A: no family; B: family; BS = breeding system: A: bi-parental care B: uni-parental care; BM = body mass; NE = nest exposure: ordinal scale of increasing nest exposure; MD = migration distance: A: resident or short migration; B: long-distance migration; BRCC = Breeding range cell account; EDGE = Evolutionary distinctiveness and globally endangered status; SW = the Shannon Wiener index for diet diversity; FSD = Foraging habitat; BSD = Body size dimorphism, A: monomorphic; B: dimorphic; PCD = Plumage color dimorphism, A: monomorphic; B: dimorphic; 1A_ ω represents ancestral node of ω for *TLR1A*. The sample coding method is applied for *TLR3*, *TLR4*, *TLR5* and *TLR7*.

| **species** | 1A_ ω | 3_ ω | 4_ ω | 5_ ω | 7_ ω | CM | FS | BS | BM | NE | MD | BRCC | EDGE | SW | FSD | BSD | PCD |
|---|---|---|---|---|---|---|---|---|---|---|---|---|---|---|---|---|---|
| ***Accipiter cooperii*** | 0.33 | / | 0.33 | 0.68 | / | B | A | B | 386.5 | B | B | 867 | 2.13 | 0.47 | 2.12 | B | A |
| ***Accipiter virgatus*** | 0.23 | 0.55 | 0.36 | / | 0.35 | B | A | B | 111.5 | A | A | 615 | 1.84 | 1.37 | 1.49 | B | A |
| ***Amazona albifrons*** | 0.27 | / | / | / | / | B | B | B | 209 | A | A | 90 | 1.63 | 1.49 | 1 | A | B |
| ***Anas platyrhynchos*** | 0.46 | 0.25 | 0.41 | 0.5 | 0.54 | A | A | A | 1162.5 | B | B | 4330 | 0.78 | 2.12 | 1.37 | A | B |
| ***Apaloderma vittatum*** | / | 0.55 | 0.32 | / | / | B | A | B | 55 | A | A | 134 | 2.8 | 0 | 1.36 | A | B |
| ***Aptenodytes forsteri*** | 0.4 | 0.21 | 0.51 | 0.35 | 0.43 | B | A | B | 32500 | B | B | 138 | 3.6 | 0.47 | 0 | A | A |
| ***Apus affinis*** | 0.42 | 0.43 | 0.38 | / | / | B | B | B | 25 | A | B | 2014 | 1.8 | 0 | 0.47 | A | A |
| ***Apus apus*** | / | 0.44 | 0.43 | / | / | B | A | B | 44 | A | B | 2356 | 2.04 | 0 | 0 | A | A |
| ***Ardeola bacchus*** | 0.37 | 0.4 | 0.56 | 0.31 | 0.37 | B | A | B | 320.4 | B | A | 385 | 2.52 | 1.16 | 1.16 | A | A |
| ***Arenaria interpres*** | 0.22 | 0.42 | 0.31 | / | 0.36 | A | A | B | 137 | B | B | 538 | 2.93 | 0.72 | 0 | A | B |
| ***Balearica regulorum*** | / | 0.24 | 0.44 | 0.35 | 0.41 | A | A | B | 3500 | B | A | 322 | 4.65 | 1.92 | 1 | A | A |
| ***Bubulcus ibis*** | 0.52 | 0.41 | 0.45 | 0.38 | 0.41 | B | A | B | 365 | B | B | 2933 | 2.92 | 1.77 | 0.88 | A | A |
| ***Buceros rhinoceros*** | / | 0.32 | 0.49 | 0.4 | 0.55 | B | B | B | 2380 | A | A | 166 | 3.36 | 1.36 | 1.16 | B | A |
| ***Bulweria bulwerii*** | 0.44 | 0.5 | 0.39 | / | / | B | A | B | 104 | A | B | 90 | 2.98 | 1 | 0 | A | A |
| ***Butorides striatus*** | 0.34 | 0.47 | 0.41 | 0.31 | 0.38 | B | B | B | 192.5 | B | A | 5076 | 2.8 | 1.57 | 0.72 | A | A |

| | | | | | | | | | | | | | | | | | |
|---|---|---|---|---|---|---|---|---|---|---|---|---|---|---|---|---|---|
| ***Calidris acuminata*** | 0.24 | 0.47 | 0.34 | / | 0.29 | A | A | A | 77.8 | B | B | 49 | 2.35 | 0.72 | 0.72 | B | A |
| ***Callaeas wilsoni*** | 0.37 | 0.18 | / | 0.36 | / | B | A | B | 226.7 | B | A | 6 | 5.28 | 1.72 | 1.57 | B | A |
| ***Caprimulgus carolinensis*** | / | 0.57 | 0.35 | 0.4 | 0.42 | A | A | B | 115.8 | B | B | 334 | 2.31 | 0.47 | 2.12 | A | B |
| ***Caprimulgus indicus*** | 0.3 | 0.38 | 0.31 | 0.37 | 0.32 | A | A | B | 81.3 | B | B | 802 | 2.6 | 0 | 1 | A | B |
| ***Cariama cristata*** | / | 0.35 | 0.39 | / | 0.58 | B | B | B | 1500 | B | A | 526 | 3.7 | 1.77 | 0.97 | A | A |
| ***Carpodacus mexicanus*** | 0.35 | 0.24 | 0.52 | / | / | B | A | B | 21.4 | B | B | 491 | 2.13 | 1.9 | 1.58 | A | B |
| ***Centropus bengalensis*** | 0.33 | 0.25 | / | 0.53 | 0.37 | B | A | B | 165 | B | A | 849 | 2.33 | 0.92 | 0.47 | B | A |
| ***Centropus sinensis*** | 0.38 | 0.25 | / | 0.5 | 0.34 | B | A | B | 252 | B | A | 781 | 2.56 | 1.97 | 0.88 | A | A |
| ***Chaetura pelagica*** | 0.42 | 0.48 | 0.33 | / | 0.36 | B | B | B | 21 | A | B | 675 | 3.02 | 0 | 1.3 | A | A |
| ***Charadrius vociferus*** | 0.38 | 0.36 | 0.4 | 0.46 | 0.27 | A | A | B | 82.5 | B | B | 1463 | 2.95 | 0.47 | 0 | A | B |
| ***Chlamydotis macqueenii*** | 0.26 | 0.32 | 0.4 | 0.4 | 0.43 | A | B | A | 1975 | B | B | 1064 | 4.55 | 2.25 | 0.72 | B | B |
| ***Clamator coromandus*** | 0.38 | 0.34 | 0.3 | 0.41 | 0.32 | B | A | B | 77 | B | B | 514 | 3.13 | 0 | 1.92 | A | A |
| ***Colius striatus*** | / | 0.37 | 0.32 | 0.39 | 0.31 | B | B | B | 51.1 | B | A | 776 | 3.54 | 1.16 | 1.76 | A | A |
| ***Columba livia*** | / | 0.25 | 0.34 | 0.46 | 0.33 | B | A | B | 267.5 | B | B | 3117 | 1.71 | 1.3 | 0.72 | A | B |
| ***Columba rupestris*** | 0.4 | 0.27 | 0.32 | 0.33 | 0.42 | B | B | B | 247.2 | A | A | 1277 | 1.71 | 1.16 | 0.72 | A | B |
| ***Cuculus canorus*** | 0.37 | 0.35 | 0.27 | 0.36 | 0.37 | B | A | B | 115 | B | B | 3301 | 1.83 | 0.47 | 1.92 | A | B |
| ***Cuculus micropterus*** | / | 0.37 | 0.29 | 0.33 | / | B | A | B | 119 | B | B | 1084 | 2.21 | 0.72 | 1.37 | A | B |
| ***Cuculus sparverioides*** | / | / | 0.26 | 0.32 | / | B | A | B | 150 | B | B | 662 | 2.37 | 0.47 | 0.97 | A | A |
| ***Cyanoramphus novaezelandiae*** | 0.3 | 0.33 | 0.21 | / | / | B | B | B | 81.5 | A | B | 33 | 2.55 | 1.9 | 1.52 | A | A |
| ***Dromaius novaehollandia*** | 0.37 | 0.23 | / | / | / | A | B | A | 31160 | B | B | 416 | 3.39 | 1.97 | 0.72 | A | B |
| ***Dromas ardeola*** | / | 0.4 | 0.37 | / | 0.21 | B | A | B | 277.5 | A | B | 229 | 3.18 | 0 | 1 | A | A |
| ***Egretta garzetta*** | 0.36 | 0.46 | 0.45 | 0.33 | 0.46 | B | A | B | 459 | B | B | 1835 | 2.19 | 1.37 | 0.88 | A | A |
| ***Elanus caeruleus*** | 0.23 | 0.57 | 0.39 | / | 0.28 | B | A | B | 259 | B | B | 2583 | 2.85 | 1.37 | 0 | B | A |
| ***Eurypyga helias*** | / | 0.32 | 0.35 | 0.23 | 0.31 | B | A | B | 188.5 | B | A | 784 | 4.03 | 1.57 | 1 | A | A |
| ***Eurystomus orientalis*** | 0.58 | / | 0.42 | / | 0.61 | B | A | B | 146.8 | A | B | 1509 | 2.65 | 0 | 2.12 | B | A |

| | | | | | | | | | | | | | | | | | |
|---|---|---|---|---|---|---|---|---|---|---|---|---|---|---|---|---|---|
| ***Falco naumanni*** | 0.25 | 0.42 | 0.39 | 0.46 | / | B | A | B | 152 | A | B | 988 | 2.26 | 0 | 1.92 | B | B |
| ***Falco peregrinus*** | 0.26 | 0.31 | 0.41 | 0.54 | 0.25 | B | B | B | 1025 | B | B | 6250 | 1.6 | 0.92 | 2.32 | B | B |
| ***Fulmarus glacialis*** | / | 0.38 | 0.39 | 0.34 | 0.42 | B | A | B | 767.5 | B | B | 443 | 2.61 | 0.97 | 0.72 | B | A |
| ***Gallicrex cinerea*** | / | / | 0.4 | 0.59 | 0.44 | A | A | B | 396 | B | B | 997 | 2.34 | 1.57 | 1.37 | B | B |
| ***Gallirallus striatus*** | / | / | 0.41 | 0.69 | 0.42 | A | A | B | 121 | B | A | 781 | 1.89 | 1.57 | 0.72 | A | A |
| ***Gallus gallus*** | 0.46 | 0.24 | 0.53 | 0.39 | 0.38 | A | A | A | 914.3 | B | A | 399 | 1.75 | 1.97 | 0 | B | B |
| ***Gavia stellata*** | / | 0.44 | / | 0.71 | / | A | A | B | 1724 | B | B | 1971 | 3.92 | 0.47 | 0 | A | A |
| ***Glareola nordmanni*** | 0.39 | 0.57 | 0.43 | / | / | A | A | B | 94.5 | B | B | 243 | 2.4 | 0 | 1.58 | A | A |
| ***Grus leucogeranus*** | 0.33 | / | 0.44 | 0.51 | 0.42 | A | B | B | 6083 | B | B | 62 | 5.16 | 2.16 | 1 | A | A |
| ***Grus nigricollis*** | 0.28 | / | 0.46 | 0.44 | / | A | B | B | 6000 | B | B | 118 | 3.26 | 2.52 | 1 | A | A |
| ***Haliaeetus albicilla*** | / | / | 0.37 | 0.76 | / | B | A | B | 4800 | B | B | 2167 | 2.1 | 1.57 | 1.76 | B | A |
| ***Haliaeetus leucocephalus*** | 0.38 | 0.53 | 0.36 | 0.82 | 0.41 | B | A | B | 4650 | B | B | 977 | 2.1 | 1.97 | 2.05 | B | A |
| ***Harpactes erythrocephalus*** | 0.28 | 0.45 | 0.36 | / | / | B | A | B | 92.5 | A | A | 310 | 2.56 | 1.36 | 1.3 | A | B |
| ***Hirundapus caudacutus*** | 0.39 | 0.39 | 0.42 | / | / | B | A | B | 120.5 | A | B | 726 | 2.78 | 0 | 0 | A | A |
| ***Ixobrychus eurhythmus*** | 0.43 | 0.42 | 0.5 | / | 0.39 | B | A | B | 142.3 | B | B | 463 | 2.95 | 1.57 | 0.88 | A | B |
| ***Ixobrychus flavicollis*** | 0.49 | 0.56 | 0.48 | / | / | B | A | B | 360 | B | B | 975 | 3.17 | 1.49 | 1 | A | B |
| ***Ixobrychus sinensis*** | 0.39 | 0.49 | 0.5 | / | 0.41 | B | A | B | 50 | B | B | 945 | 2.96 | 0.92 | 1 | A | B |
| ***Larus mongolicus*** | 0.41 | 0.49 | 0.38 | 0.45 | 0.36 | A | B | B | 1150 | B | B | 482 | 0.74 | 1.85 | 1.57 | A | A |
| ***Leptosomus discolor*** | / | 0.21 | 0.36 | 0.35 | 0.46 | B | A | B | 231 | A | A | 58 | 4.29 | 0.88 | 1.52 | A | B |
| ***Meleagris gallopavo*** | / | 0.29 | 0.48 | / | 0.35 | A | B | A | 5810 | B | A | 557 | 2.32 | 1.92 | 0.47 | B | B |
| ***Melopsittacus undulatus*** | 0.34 | 0.27 | 0.25 | 0.44 | 0.35 | B | A | B | 27.5 | A | B | 561 | 2.65 | 0 | 0.97 | A | B |
| ***Merops nubicus*** | / | 0.44 | 0.35 | 0.6 | 0.64 | B | B | B | 52.5 | A | B | 641 | 2.25 | 0 | 1.96 | A | A |
| ***Merops viridis*** | 0.53 | 0.38 | 0.35 | / | 0.38 | B | B | B | 37.5 | A | B | 407 | 1.98 | 0 | 1.3 | A | A |
| ***Mesitornis unicolor*** | / | 0.41 | 0.48 | 0.36 | / | A | A | B | 148 | B | A | 37 | 4.77 | 0 | 0 | A | A |
| ***Mesophoyx intermedia*** | 0.51 | 0.52 | 0.51 | 0.34 | 0.42 | B | A | B | 400 | B | A | 2237 | 2.77 | 1.16 | 0.72 | A | A |

| | | | | | | | | | | | | | | | | | |
|---|---|---|---|---|---|---|---|---|---|---|---|---|---|---|---|---|---|
| ***Mohoua ochrocephala*** | 0.38 | 0.33 | / | 0.37 | / | B | B | B | 30 | A | A | 12 | 4.71 | 1.37 | 1 | A | A |
| ***Nestor notabilis*** | / | 0.31 | 0.34 | / | 0.22 | B | A | B | 922 | A | B | 13 | 4.4 | 1.3 | 1.85 | A | B |
| ***Nipponia nippon*** | 0.43 | 0.24 | 0.38 | 0.36 | 0.38 | B | B | B | 1900 | B | A | 7 | 5.34 | 1.16 | 0.72 | A | A |
| ***Notiomystis cincta*** | 0.3 | / | 0.56 | / | / | B | A | B | 93.9 | A | A | 3 | 4.86 | 1.57 | 1.37 | B | B |
| ***Nycticorax nycticorax*** | 0.43 | 0.46 | 0.42 | 0.31 | 0.49 | B | A | B | 662.5 | B | B | 3975 | 3.05 | 1.97 | 1 | A | A |
| ***Nyctyornis athertoni*** | / | 0.44 | 0.34 | / | 0.45 | B | A | B | 84.8 | A | A | 267 | 3.33 | 0 | 0.97 | B | A |
| ***Oceabdroma leucorhoa*** | 0.35 | 0.43 | 0.43 | 0.39 | / | B | A | B | 45 | A | B | 18 | 3.09 | 1.49 | 0.72 | A | A |
| ***Ophisthocomus hoazin*** | 0.54 | 0.31 | 0.25 | 0.3 | 0.51 | B | B | B | 800 | B | A | 615 | 4.23 | 0 | 0.72 | A | A |
| ***Otis tarda*** | 0.27 | 0.21 | 0.47 | 0.38 | 0.43 | A | B | A | 8100 | B | B | 477 | 4.55 | 1.72 | 0.72 | B | B |
| ***Otus sunia*** | 0.35 | 0.38 | 0.35 | / | 0.42 | B | A | B | 105 | A | B | 327 | 2.53 | 1.16 | 1.96 | A | B |
| ***Otus spilocephalus*** | / | / | 0.35 | / | 0.44 | B | A | B | 87.5 | A | A | 391 | 2.4 | 0.72 | 1.36 | A | A |
| ***Pelecanus crispus*** | / | 0.55 | 0.51 | 0.63 | 0.65 | B | A | B | 11500 | B | B | 224 | 4.57 | 0 | 0 | A | A |
| ***Pelecanus rufescens*** | 0.43 | 0.49 | / | / | / | B | A | B | 5450 | B | B | 1616 | 3.16 | 0 | 0 | A | A |
| ***Perdix perdix*** | / | / | 0.5 | 0.43 | 0.3 | A | B | B | 380 | B | A | 955 | 2 | 1.57 | 0 | B | B |
| ***Petroica australis*** | 0.33 | 0.25 | / | 0.42 | / | B | A | B | 40 | B | A | 21 | 2.62 | 0.72 | 0.47 | A | A |
| ***Phaethon lepturus*** | / | 0.25 | 0.46 | 0.25 | 0.27 | B | A | B | 315 | B | B | 283 | 3.51 | 0.88 | 0.72 | A | A |
| ***Phalacrocorax carbo*** | / | 0.21 | / | 0.38 | / | B | A | B | 2360 | B | B | 1738 | 2.2 | 0.92 | 0 | A | A |
| ***Philesturnus carunculatus*** | 0.39 | / | / | 0.38 | / | B | A | B | 75 | A | A | 2 | 3.9 | 1.36 | 1.92 | B | A |
| ***Phoenicopterus roseus*** | 0.53 | 0.82 | 0.4 | 0.42 | 0.37 | B | A | B | 3031.6 | A | A | 342 | 3.04 | 1.76 | 0 | B | A |
| ***Picoides pubescens*** | 0.27 | 0.31 | 0.3 | 0.48 | / | B | A | B | 26.5 | A | B | 1197 | 1.8 | 0.92 | 1.52 | A | B |
| ***Podiceps cristatus*** | 0.35 | / | 0.41 | / | 0.34 | A | B | B | 1043 | B | B | 2226 | 2.8 | 1.3 | 0.88 | A | A |
| ***Porphyrio hochstetteri*** | 0.3 | 0.39 | 0.38 | / | 0.38 | A | B | B | 2470 | B | A | 1 | 4.67 | 1.69 | 0.97 | A | A |
| ***Pterocles gutturalis*** | / | 0.51 | 0.35 | 0.44 | 0.19 | A | A | B | 342.5 | B | B | 199 | 3.04 | 0 | 0.88 | B | B |
| ***Pygoscelis adeliae*** | / | 0.27 | 0.46 | 0.33 | / | B | A | B | 4880 | B | B | 138 | 3.83 | 0.88 | 0 | A | A |
| ***Streptopelia chinensis*** | 0.53 | 0.24 | 0.34 | / | / | B | A | B | 128 | B | B | 916 | 2.53 | 0 | 0 | A | A |

| | | | | | | | | | | | | | | | | | |
|---|---|---|---|---|---|---|---|---|---|---|---|---|---|---|---|---|---|
| ***Streptopelia tranquebarica*** | 0.45 | 0.24 | 0.32 | 0.46 | / | B | A | B | 104 | B | B | 874 | 2.16 | 0.97 | 0 | A | B |
| ***Strigops habroptila*** | 0.35 | 0.39 | 0.26 | / | / | B | B | A | 1975 | A | A | 2 | 6.28 | 1.49 | 0 | A | A |
| ***Struthio camelus*** | 0.35 | 0.19 | 0.37 | 0.66 | 0.25 | A | B | B | 116500 | B | B | 919 | 4.02 | 1.69 | 0 | B | B |
| ***Taeniopygia guttata*** | 0.42 | 0.24 | 0.57 | 0.38 | / | B | A | B | 12 | A | A | 43 | 2.11 | 0 | 0.72 | A | B |
| ***Tauraco erythrolophus*** | / | 0.42 | 0.38 | 0.26 | 0.39 | B | A | B | 267.5 | B | A | 52 | 3.08 | 0.97 | 1.49 | A | A |
| ***Tinamus guttatus*** | / | 0.22 | 0.51 | 0.4 | 0.28 | A | B | A | 688.8 | B | A | 381 | 3.88 | 1.37 | 0 | B | A |
| ***Tringa totanus*** | 0.18 | 0.39 | 0.28 | / | / | A | A | B | 120 | B | B | 1560 | 2.5 | 0.92 | 0.72 | A | A |
| ***Turnix suscitator*** | 0.33 | / | 0.35 | / | 0.28 | A | A | A | 50.5 | B | A | 735 | 2.44 | 1.49 | 0 | B | B |
| ***Turnix tanki*** | 0.32 | / | 0.33 | / | 0.28 | A | A | A | 79.8 | B | B | 759 | 2.44 | 1.37 | 0 | B | B |
| ***Tyto alba*** | / | 0.39 | 0.38 | 0.22 | 0.23 | B | A | B | 321 | A | A | 6303 | 2.72 | 0.92 | 0.47 | B | A |
| ***Urocissa erythrorhyncha*** | 0.35 | 0.29 | 0.5 | / | / | B | B | B | 160.4 | B | A | 484 | 1.98 | 2.12 | 1.49 | B | A |
| ***Xenicus gilviventris*** | 0.37 | 0.34 | / | / | / | B | A | B | 17.8 | A | A | 9 | 4.97 | 0.92 | 1 | B | B |
| ***Zoothera dauma*** | 0.39 | / | 0.63 | 0.44 | 0.35 | B | A | B | 95.5 | B | B | 911 | 2.14 | 0.72 | 0.97 | A | A |

**Table S7** Phylogenetic generalized linear mixed models using MCMCglmm (Hadfield 2010) comparing the effect of life-history and ecological variables on the evolution of ω of different *TLRs*. Model selection and model averaging approach according to the DICC (ΔDICc < 2). Factors with a sum of Akaike weights (Σ AICC weights) larger than 0.5 and the 95% CI of estimates do not overlap 0 are highlighted in bold.

| TLR number / parameter | Estimate | SE | L 95% CI | U 95% CI | Σ DIC weights | N models incl. variable |
|---|---|---|---|---|---|---|
| *TLR1LA* | | | | | | |
| **Intercept** | **0.336** | **0.07** | **0.202** | **0.471** | | |
| Log (body mass) | 0.01 | 0.01 | -0.003 | 0.023 | 1 | 44 |
| **development mode (precocial vs altricial)** | **0.068** | **0.03** | **0.002** | **0.134** | **0.96** | **42** |
| plumage color dimorphism (mono- vs di-morphism) | -0.027 | 0.02 | -0.063 | 0.01 | 0.91 | 41 |
| **diet diversity** | **-0.045** | **0.02** | **-0.081** | **-0.009** | **0.73** | **31** |
| family living (non-family vs family living) | -0.018 | 0.02 | -0.061 | 0.025 | 0.52 | 23 |
| body size dimorphism (mono- vs di-morphism) | -0.017 | 0.02 | -0.061 | 0.027 | 0.48 | 21 |
| Log (breeding range cell account) | 0.003 | 0.01 | -0.007 | 0.013 | 0.26 | 11 |
| breeding system (biparental vs uniparental care) | -0.029 | 0.04 | -0.098 | 0.041 | 0.24 | 11 |
| EDGE | 0.003 | 0.01 | -0.016 | 0.023 | 0.17 | 8 |
| sedentariness (resident vs migratory) | 0.009 | 0.02 | -0.026 | 0.044 | 0.16 | 8 |
| foraging habitat | -0.005 | 0.02 | -0.041 | 0.03 | 0.16 | 7 |
| nest exposure (closed vs open) | 0.008 | 0.02 | -0.036 | 0.052 | 0.14 | 7 |
| *TLR3* | | | | | | |
| **Intercept** | **-0.897** | **0.26** | **-1.398** | **-0.396** | | |
| Log (body mass) | -0.035 | 0.02 | -0.076 | 0.005 | 1 | 5 |
| development mode (precocial vs altricial) | -0.102 | 0.1 | -0.29 | 0.085 | 1 | 5 |
| EDGE | -0.556 | 0.54 | -1.616 | 0.504 | 0.51 | 2 |
| Log (breeding range cell account) | -0.202 | 0.2 | -0.589 | 0.185 | 0.51 | 2 |
| breeding system (biparental vs uniparental care) | 0.045 | 0.13 | -0.204 | 0.293 | 0.39 | 2 |

| TLR number / parameter | Estimate | SE | L 95% CI | U 95% CI | Σ DIC weights | N models incl. variable |
|---|---|---|---|---|---|---|
| foraging habitat | 0.013 | 0.06 | -0.114 | 0.139 | 0.27 | 2 |
| sedentariness (resident vs migratory) | -0.018 | 0.05 | -0.119 | 0.082 | 0.15 | 1 |
| family living (non-family vs family living) | 0.021 | 0.05 | -0.08 | 0.122 | 0.12 | 1 |
| plumage color dimorphism (mono- vs di-morphism) | -0.077 | 0.07 | -0.212 | 0.058 | 0.12 | 1 |
| diet diversity | -0.036 | 0.06 | -0.161 | 0.088 | 0.12 | 1 |
| *TLR4* | | | | | | |
| **Intercept** | **0.429** | **0.06** | **0.303** | **0.555** | | |
| **family living (non-family vs family living)** | **-0.033** | **0.01** | **-0.059** | **-0.007** | **1** | **41** |
| **Log (body mass)** | **0.012** | **0.01** | **0.002** | **0.022** | **1** | **41** |
| EDGE | -0.012 | 0.01 | -0.03 | 0.006 | 0.94 | 39 |
| sedentariness (resident vs migratory) | -0.016 | 0.01 | -0.04 | 0.008 | 0.85 | 34 |
| Log (breeding range cell account) | -0.006 | 0.01 | -0.017 | 0.004 | 0.76 | 30 |
| development mode (precocial vs altricial) | -0.022 | 0.02 | -0.066 | 0.022 | 0.74 | 30 |
| diet diversity | -0.004 | 0.01 | -0.027 | 0.018 | 0.39 | 15 |
| foraging habitat | -0.014 | 0.01 | -0.037 | 0.01 | 0.39 | 18 |
| body size dimorphism (mono- vs di-morphism) | 0.012 | 0.02 | -0.022 | 0.046 | 0.26 | 12 |
| breeding system (biparental vs uniparental care) | 0.007 | 0.02 | -0.035 | 0.048 | 0.2 | 9 |
| nest exposure (closed vs open) | 0.016 | 0.02 | -0.017 | 0.048 | 0.13 | 6 |
| plumage color dimorphism (mono- vs di-morphism) | -0.007 | 0.02 | -0.037 | 0.023 | 0.04 | 1 |
| *TLR5* | | | | | | |
| **Intercept** | **-1.013** | **0.17** | **-1.346** | **-0.68** | | |
| **sedentariness (resident vs migratory)** | **0.195** | **0.06** | **0.08** | **0.311** | **1** | **6** |
| nest exposure (closed vs open) | 0.072 | 0.06 | -0.05 | 0.194 | 1 | 6 |
| foraging habitat | 0.013 | 0.05 | -0.089 | 0.114 | 0.55 | 3 |
| Log (breeding range cell account) | -0.002 | 0.02 | -0.041 | 0.038 | 0.43 | 2 |

| TLR number / parameter | Estimate | SE | L 95% CI | U 95% CI | Σ DIC weights | N models incl. variable |
|---|---|---|---|---|---|---|
| breeding system (biparental vs uniparental care) | 0.072 | 0.13 | -0.175 | 0.319 | 0.39 | 3 |
| diet diversity | -0.004 | 0.05 | -0.097 | 0.088 | 0.33 | 2 |
| plumage color dimorphism (mono- vs di-morphism) | 0.013 | 0.07 | -0.133 | 0.158 | 0.16 | 1 |
| *TLR7* | | | | | | |
| **Intercept** | **0.189** | **0.1** | **-0.012** | **0.39** | | |
| **Log (body mass)** | **0.018** | **0.01** | **0.003** | **0.033** | **1** | **23** |
| Log (breeding range cell account) | 0.009 | 0.01 | -0.006 | 0.025 | 1 | 23 |
| foraging habitat | 0.039 | 0.02 | -0.001 | 0.078 | 1 | 23 |
| nest exposure (closed vs open) | -0.035 | 0.03 | -0.096 | 0.026 | 0.85 | 20 |
| sedentariness (resident vs migratory) | -0.019 | 0.02 | -0.065 | 0.028 | 0.55 | 12 |
| development mode (precocial vs altricial) | 0.002 | 0.04 | -0.068 | 0.071 | 0.32 | 9 |
| breeding system (biparental vs uniparental care) | -0.025 | 0.04 | -0.103 | 0.053 | 0.27 | 5 |
| EDGE | 0.002 | 0.02 | -0.032 | 0.036 | 0.21 | 8 |
| diet diversity | -0.007 | 0.02 | -0.044 | 0.029 | 0.17 | 5 |
| family living (non-family vs family living) | 0.007 | 0.03 | -0.043 | 0.057 | 0.14 | 3 |
| plumage color dimorphism (mono- vs di-morphism) | -0.028 | 0.03 | -0.085 | 0.029 | 0.09 | 3 |
